# Pathogen and host biomarkers to aid early diagnosis and prognosis of tuberculous meningitis

**DOI:** 10.64898/2026.05.27.728147

**Authors:** Urvashi B Singh, Angitha K P, Adarsh A K, Kiran Singh, Naveet Wig, Achal Kumar Srivastava, Uma Kanga

## Abstract

**Background:** Tuberculous meningitis (TBM) is the most sinister form of extrapulmonary tuberculosis (EPTB), associated with high mortality due to delayed diagnosis and limited sensitivity of conventional and molecular tests. Current study evaluated the diagnostic utility of Lipoarabinomannan antigen (LAM) detection in CSF and urine and explored host inflammatory biomarkers for diagnosis and prognosis of TBM.

**Methods:** This prospective observational study enrolled 80 patients with presumptive TBM at a tertiary care centre. CSF samples were subjected to AFB microscopy, liquid culture(MGIT-960), GeneXpert MTB/RIF (GX), and LAM lateral flow assay. Urine LAM was performed at baseline. Serum and CSF levels of IL-1β, IL-6, TNF-α, IFN-γ, IL-17A, and IP-10 were measured at baseline and after 1 month treatment.

**Results:** Among 80 participants, 23 (28.7%) had definite TBM and 46 (57.5%) had probable TBM. CSF LAM sensitivity and specificity against microbiological reference standards was 43.5% and 80.7%, while urine LAM sensitivity (60.9%) and specificity 82.5% was higher. Against composite reference standards, both CSF and urine LAM showed reduced sensitivity but achieved 100% specificity. Serum IL-1β showed the best diagnostic performance (AUC 0.943; sensitivity 88.9%, specificity 90.9%). Elevated serum and CSF IP-10 levels were associated with poor outcomes, whereas declining IL-6 and TNF-α levels correlated with treatment response.

**Conclusion:** LAM detection in CSF and urine may serve as a highly specific, rapid rule-in test for TBM. Host inflammatory biomarkers, especially IL-1β and IP-10, show additional diagnostic and prognostic value. Combining LAM testing with cytokine biomarkers may improve early diagnosis and efficient clinical management of TBM.

## 1. Introduction

*Mycobacterium tuberculosis* (MTB) affects 9.9 million people worldwide and carries devastating consequences (1). Although lungs are the primary site of TB infection, EPTB accounts for 15% of reported cases (2). TBM accounts for 1% of all cases of active tuberculosis and 5% to 10% of cases of EPTB (3). Central nervous system tuberculosis (CNS TB) is challenging to diagnose and has a poor prognosis, if treatment is delayed (4).

The most common rapid diagnostic methods include cerebrospinal fluid (CSF) smear microscopy and molecular tests, such as the GeneXpert MTB/RIF assay [(Cepheid, Sunnyvale, CA, USA) (GX)] (4). However, microscopic detection is often difficult due to CSF low bacillary load (6), whereas gold-standard CSF culture has only moderate sensitivity (30–60%) (3). Systematic reviews and meta-analyses evaluating GX for TBM have reported sensitivities ranging from 62.8% to 66% and specificities between 89% to 98.8% against composite reference standards (CRS) (5). However, its reduced specificity and positive predictive value (PPV) necessitate further prospective evaluation in larger and more diverse populations, such as children and HIV-uninfected individuals (6).

There is an urgent need for a rapid biomarker-based test with high sensitivity and low cost (7). Lipoarabinomannan, a cell wall component of MTB, which is excreted in urine, is a promising biomarker (8). Currently, the WHO recommends the lateral flow LAM assay (LFA) in urine for diagnosis of TB in PLHIV (7). Although CSF-based biomarkers have shown potential for TBM diagnosis (9), CSF collection requires an invasive lumbar puncture (9). As a result, blood-based host protein biomarkers have emerged as promising alternatives, with recent studies evaluating their potential in pediatric TBM (9).

The current study was designed to look for the diagnostic utility of LAM antigen detection in CSF and urine for the diagnosis of TBM. This study also aimed to explore the diagnostic potential of host serum and CSF biomarkers for TBM.

## 2. Materials and Methods

### 2.1 Study Design and Population

This prospective observational study was conducted in the Departments of Microbiology, Medicine, and Neurology at the All India Institute of Medical Sciences (AIIMS), New Delhi, India, from July 2020 to December 2022. During the study period, a total of 80 patients were enrolled. Patients already receiving anti-tubercular therapy, pregnant women, people living with HIV, and those who did not give informed consent were excluded. The workflow of the study is summarized in **Supplementary 1.**

### 2.2. Clinical Data Collection

A standardized proforma was used to document demographic information, presenting symptoms (fever, headache, vomiting, altered sensorium, and seizures), constitutional symptoms, previous tuberculosis treatments, and underlying comorbidities. Microbiological outcomes were documented, including microscopy, culture, GX, and detection of LAM in urine and CSF. Clinical outcomes such as improvement or death were recorded during follow-up.

### 2.3 Case definition

The consistent case definition for TBM proposed by *Marais et al*. (10) was used to classify patients as having definitive, probable, possible, or non-TBM based on clinical, CSF, radiographic, and microbiological findings.

### 2.4 Sample Collection and Processing

Clinical samples, including CSF, urine, and blood, were collected from patients with high suspicion of meningitis. AFB smear, liquid culture with MGIT 960 [(Becton Dickinson, Franklin Lakes, NJ, USA) (LC)], and GX tests were performed on CSF according to standard guidelines. (11,12). LAM detection by LFA was performed on CSF and urine, and biomarkers were assessed in CSF and serum per the manufacturer’s instructions.

#### 2.4.1 TB LAM Antigen Detection

All CSF and urine samples were tested with the Determine®-TB LAM Ag LFA (Abbott, Chicago, IL, USA). FUJILAM kits (Fujifilm, Tokyo, Japan) were unavailable in the country during the study; Abbott Laboratories’ Determine TM TB LAM Ag test kits were used. Samples were centrifuged at 10,000g for 5 minutes, and 60 µL of clear supernatant was pipetted onto strip. Results were read after 25–35 minutes.

#### 2.4.2. Serum/CSF Biomarkers

For biomarker analysis, an additional 1 mL of CSF and blood collected in a BD Vacutainer® serum tube were processed by centrifugation, aliquoted, and stored at −80°C until testing. Samples were collected at baseline (before treatment began) and at 1-month (FU-1) intervals. Cytokine levels, including CXCL10 /IP-10 (Interferon-γ inducible protein −10), interferon gamma (IFN-γ), interleukin-1-beta (IL-1β), IL-6, IL-17A, and tumor necrosis factor alpha (TNF-α) levels were measured in serum and CSF using Pro-Human Cytokine Assay (Bio-Plex; Bio-Rad Laboratories, Hercules, CA, USA) according to the manufacturer’s instructions.

### 2.5 Statistical analysis

Statistical analyses were performed using SPSS version 25.0. Continuous variables are expressed as mean (standard deviation) or median (interquartile range {IQR}), while categorical variables are reported as frequencies (n) and percentages (%). A p-value of less than 0.05 was considered statistically significant. Numeric variables were compared using a t-test. Chi-square tests examined differences in categorical variables among definite, probable, and non-TBM groups. The median cytokine levels (IQR) for the TBM and controls were compared using nonparametric Mann-Whitney U test.

Diagnostic accuracy parameters, with 95% confidence intervals (CIs), were estimated for CSF TB-LAM, urine TB-LAM, and GX assay against definite TBM (positive LC/GX) and a composite reference of probable or definite TBM, based on uniform case definition. (14) Receiver operator characteristic (ROC) curves for identified biomarkers were generated for IL-6, TNF-α, IP-10, IL-1β, IFN-γ, and IL-17A, with optimal cut-off values determined using the maximum Youden index (sensitivity + specificity − 1).

## Results

Overall, eighty hospitalized participants with presumptive TBM were enrolled in the study. The mean age of patients was 30.5 years (SD = 14.7), with males comprising 56.2% of the cohort. **Table 1** provides an overview of the study participants’ characteristics.

**Table 1.**
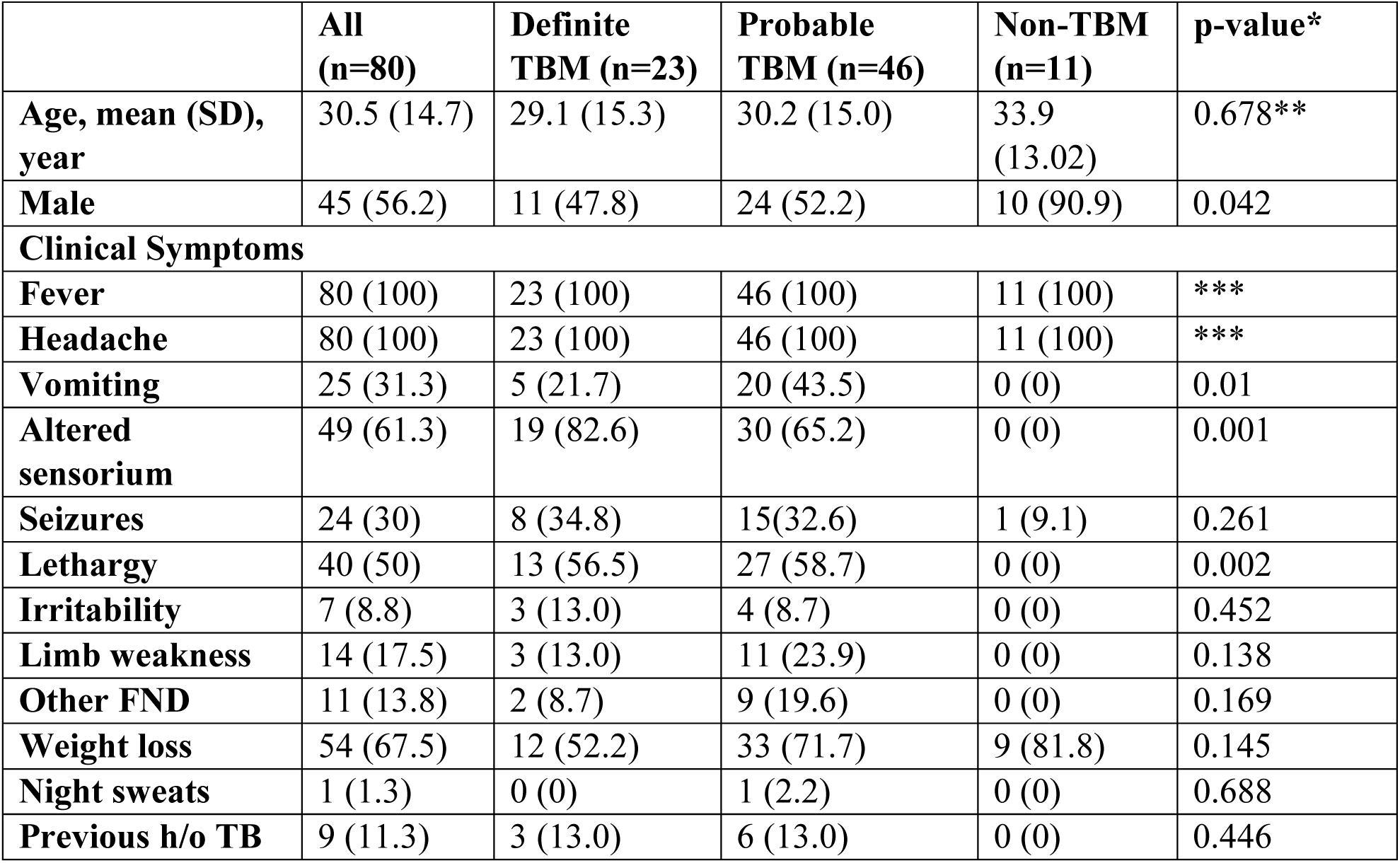
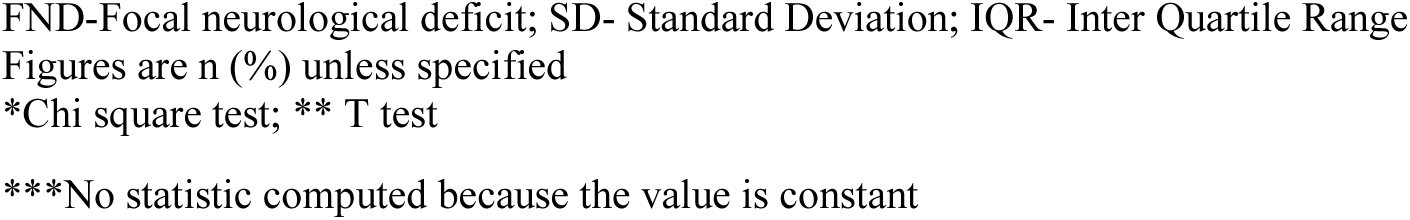
Characteristics of study participants

|  | **All**<br>**(n=80)** | **Definite**<br>**TBM (n=23)** | **Probable**<br>**TBM (n=46)** | **Non-TBM**<br>**(n=11)** | **p-value\*** |
| --- | --- | --- | --- | --- | --- |
| **Age, mean (SD),**<br>**year** | 30.5 (14.7) | 29.1 (15.3) | 30.2 (15.0) | 33.9<br>(13.02) | 0.678** |
| **Male** | 45 (56.2) | 11 (47.8) | 24 (52.2) | 10 (90.9) | 0.042 |
| **Clinical Symptoms** |  |  |  |  |  |
| **Fever** | 80 (100) | 23 (100) | 46 (100) | 11 (100) | *** |
| **Headache** | 80 (100) | 23 (100) | 46 (100) | 11 (100) | *** |
| **Vomiting** | 25 (31.3) | 5 (21.7) | 20 (43.5) | 0 (0) | 0.01 |
| **Altered**<br>**sensorium** | 49 (61.3) | 19 (82.6) | 30 (65.2) | 0 (0) | 0.001 |
| **Seizures** | 24 (30) | 8 (34.8) | 15(32.6) | 1 (9.1) | 0.261 |
| **Lethargy** | 40 (50) | 13 (56.5) | 27 (58.7) | 0 (0) | 0.002 |
| **Irritability** | 7 (8.8) | 3 (13.0) | 4 (8.7) | 0 (0) | 0.452 |
| **Limb weakness** | 14 (17.5) | 3 (13.0) | 11 (23.9) | 0 (0) | 0.138 |
| **Other FND** | 11 (13.8) | 2 (8.7) | 9 (19.6) | 0 (0) | 0.169 |
| **Weight loss** | 54 (67.5) | 12 (52.2) | 33 (71.7) | 9 (81.8) | 0.145 |
| **Night sweats** | 1 (1.3) | 0 (0) | 1 (2.2) | 0 (0) | 0.688 |
| **Previous h/o TB** | 9 (11.3) | 3 (13.0) | 6 (13.0) | 0 (0) | 0.446 |

FND-Focal neurological deficit; SD- Standard Deviation; IQR- Inter Quartile Range Figures are n (%) unless specified \*Chi square test; \*\* T test
\*\*\*No statistic computed because the value is constant

Based on the criteria of *Marais et al.* (10), definite TBM was defined as detection of MTB in CSF by smear, culture, or GX. The TBM group included both definite and probable TBM cases (10), while the non-TBM group included other meningitis etiologies, including cryptococcal meningitis (10).

Definite TBM was diagnosed in 23 patients (28.7%) based on the microbiologic reference standard (MRS) (GX, n = 19; MGIT, n = 12). Probable TBM was diagnosed in an additional 46 patients (57.5%) according to the case definitions (10). Using CRS, 69 patients (86.2%) were diagnosed as definite or probable TBM. The remaining 11 patients were classified as non-TBM and had cryptococcal meningitis (CSF cryptococcal antigen test/culture positive). **Supplementary 2** demonstrates the CT findings in the various study groups. **Supplementary 3a and 3b** detail CSF parameters based on definite/probable/non-TBM diagnosis. **Supplementary 4 and 5** demonstrate the BMRC staging and clinical outcome in patients, respectively.

Among definite TBM cases, GX identified 82.6% (19/23), whereas MGIT culture detected 52.2% (12/23). None of the CSF samples were smear positive. Four (5%) LC-positive participants were GX-negative, and eleven (13.75%) GX-positive participants were LC-negative. Fifty-seven (71.25%) participants had negative CSF LC and GX results. **Table 2** provides an overview of test outcomes.

**Table 2:** Microbiological investigations in study participants

|  | **All**<br>**(n=80)** | **Definite TBM**<br>**(n=23)** | **Probable TBM (n=46)** | **Non-TBM**<br>**(n=11)** |
| --- | --- | --- | --- | --- |
| **CSF sample** |  |  |  |  |
| **AFB** | 0(0) | 0 (0) | 0 (0) | 0 (0) |
| **LC** | 12 (15.0) | 12 (52.2) | 0 (0) | 0 (0) |
| **GX** | 19 (23.7) | 19 (82.6) | 0 (0) | 0 (0) |
| **CSF LAM** | 21 (26.2) | 10 (43.5) | 11 (23.9) | 0 (0) |
| **Urine sample** |  |  |  |  |
| **URINE LAM** | 24 (30.0) | 14 (60.8) | 10 (21.7) | 0 (0) |

AFB-acid fast bacilli; LC-Liquid Culture; GX-GeneXpert; LAM-Lipoarabinomannan Figures are n (%) unless specified.

LAM antigen was detected in CSF samples from 10 patients with definite TBM and 11 with probable TBM, while all non-TBM cases tested negative. In urine samples, LAM antigen was detected in 14 definite TBM cases and 10 probable TBM cases, with no positivity observed in the non-TBM group. **Figure 1a and 1b** demonstrate diagnostic overlap of CSF GeneXpert, MGIT culture, CSF LAM, and urine LAM positivity among confirmed TBM cases.

**Figure 1a.**
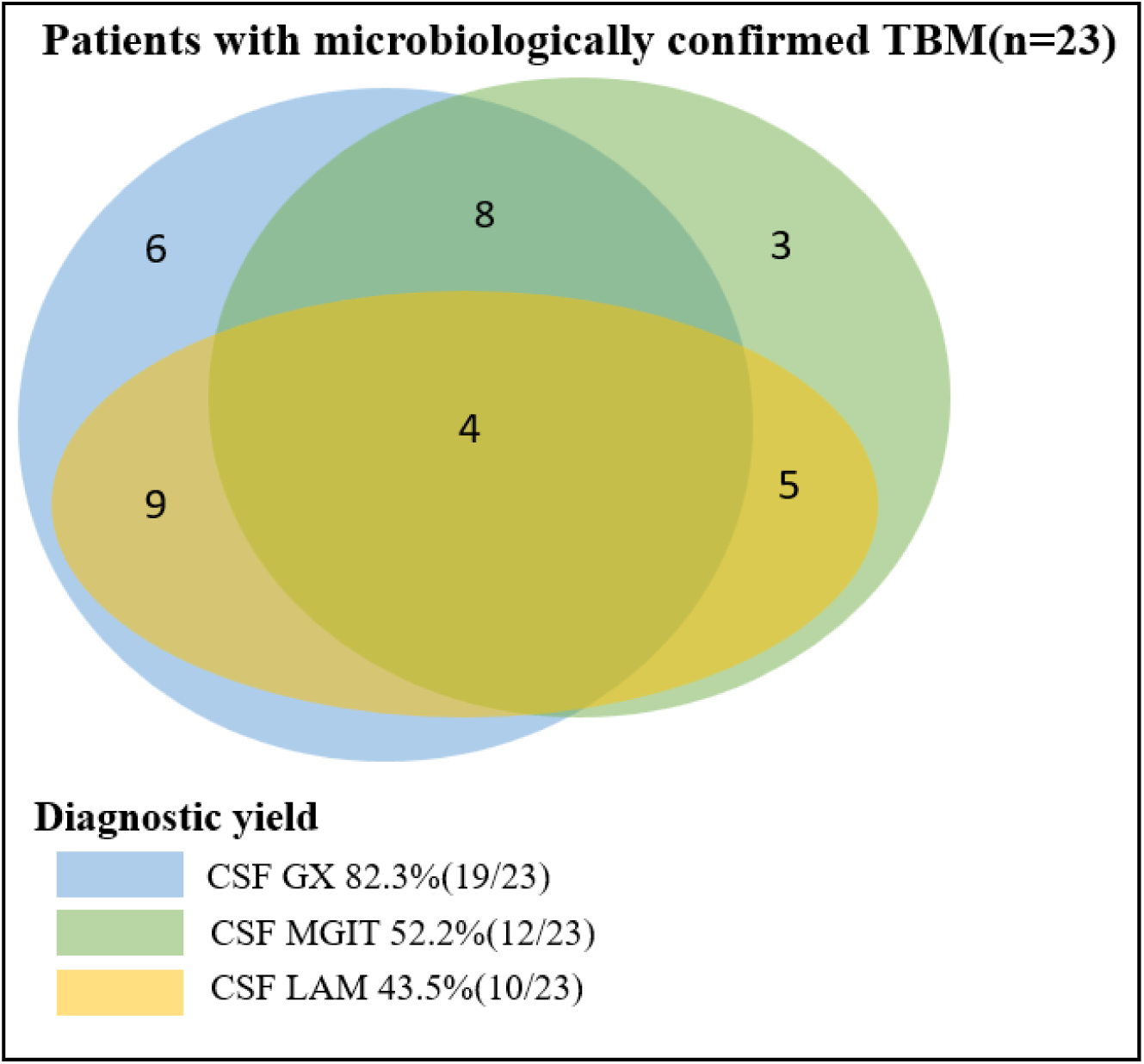
Diagnostic overlap of CSF GeneXpert, MGIT, and LAM in confirmed TBM (n=23). The The Venn diagram displays 23 participants with definite TBM detected in CSF by each diagnostic test and the overlap between tests. GX-GeneXpert; LAM- Lipoarabinomannan; MGIT- mycobacterial growth indicator tube

**Figure 1b.**
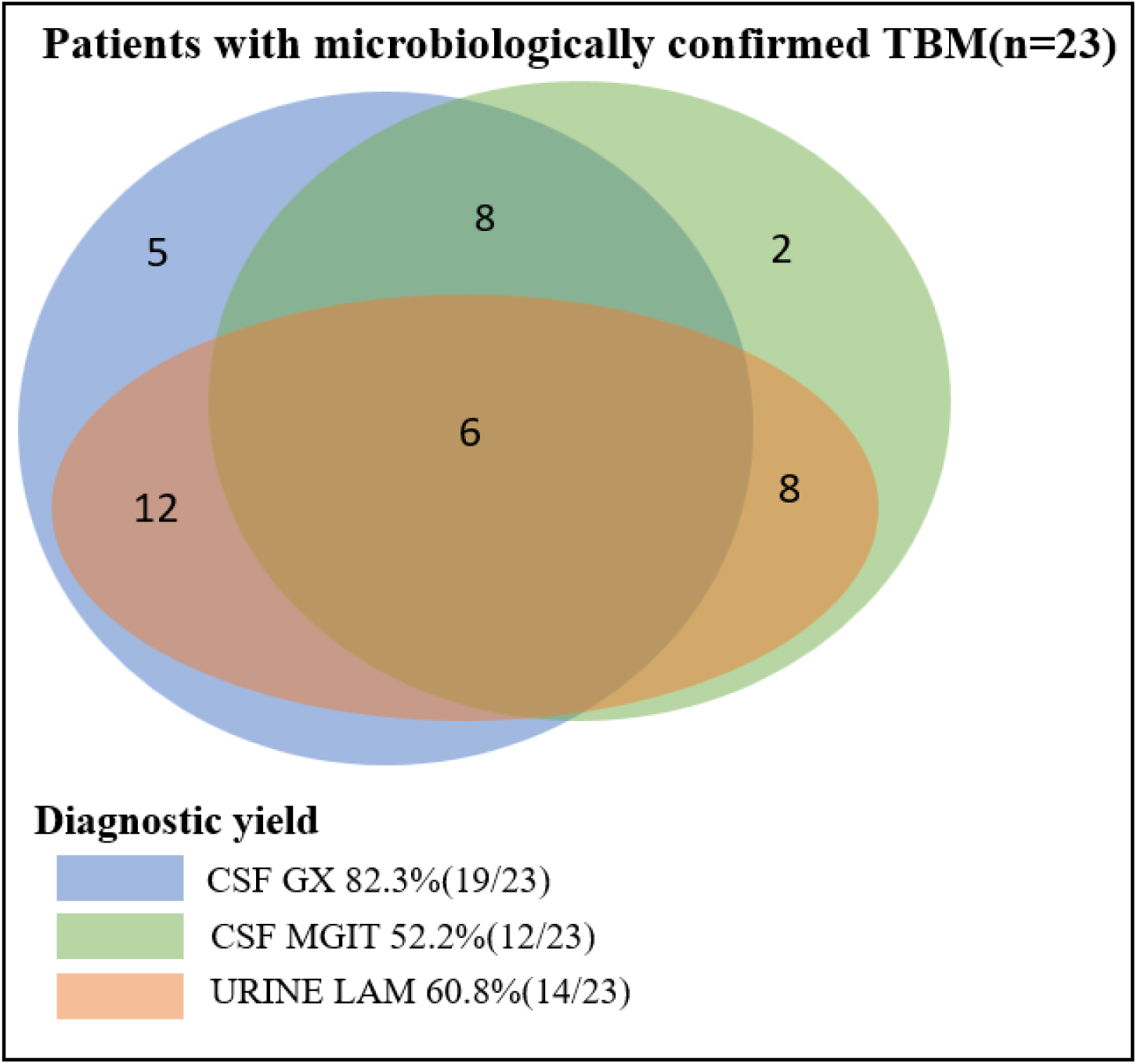
Diagnostic overlap of CSF GeneXpert, MGIT, and LAM in confirmed TBM (n=23). The The Venn diagram displays 23 participants with definite TBM detected in CSF by each diagnostic test and the overlap between tests. GX-GeneXpert; LAM- Lipoarabinomannan; MGIT- mycobacterial growth indicator tube

Against CRS, CSF LAM showed a sensitivity of 30.4% and specificity of 100%. The sensitivity of CSF LAM detection improved to 43.5%, while the specificity decreased to 80.7% against MRS. The diagnostic performance parameters for CSF LAM are presented in **Table 3**.

**Table 3.** Diagnostic performance of CSF mycobacterial LAM assay for tuberculous meningitis

| **Reference Standard** | **Sensitivity**<br>**n/N (%)** | **Specificity**<br>**n/N (%)** | **PPV**<br>**n/N (%)** | **NPV**<br>**n/N (%)** | **Positive**<br>**Likelihood**<br>**Ratio** | **Negative**<br>**Likelihood**<br>**Ratio** |
| --- | --- | --- | --- | --- | --- | --- |
| **Comprehensive Reference Standard** | 21/69<br>(30.4%) | 11/11<br>(100%) | 21/21<br>(100%) | 11/59<br>(18.6%) | - | 0.70 |
| **95% CI** | 19.9 to<br>42.7% | 71.5 to<br>100% | - | 16.4 to<br>21.1% | - | 0.6 to 0.8% |
| **Microbiological reference standard** | 10/23<br>(43.5%) | 46/57<br>(80.7%) | 10/21<br>(47.6%) | 46/59<br>(77.9%) | 2.25 | 0.70 |
| **95% CI** | 23.2 to<br>65.5% | 68.1 to<br>89.9% | 30.9 to<br>64.8% | 70.7 to<br>83.8% | 1.1 to 4.6% | 0.5 to 1.0% |
| **GX** | 9/19<br>(47.4%) | 49/61<br>(80.3%) | 9/21<br>(42.9%) | 49/59<br>(83.0%) | 2.41 | 0.66 |
| **95% CI** | 24.4 to<br>71.1% | 68.2 to<br>89.4% | 27.2 to<br>60.0% | 75.9 to<br>88.4% | 1.2 to 4.8% | 0.4 to 1.0% |
| **MGIT** | 5/12<br>(41.7%) | 52/68<br>(76.5%) | 5/21<br>(23.8%) | 52/59<br>(88.1%) | 1.7 | 0.7 |
| **95% CI** | 15.2 to<br>72.3% | 64.6 to<br>85.9% | 12.4 to<br>40.9% | 81.9 to<br>92.4% | 0.8 to 3.9% | 0.5 to 1.2% |

PPV-Positive predictive value; NPV-Negative predictive value; CI- Confidence Interval
GX-GeneXpert; MGIT- Mycobacterial Growth Indicator Tube

Urine LAM against CRS showed a sensitivity of 34.5% and specificity of 100%. It exhibited higher sensitivity than CSF LAM against MRS at 60.9%, but lower specificity at 82.5%. The sensitivity, specificity, PPV, and NPV of urine LAM against CRS, MRS, GX, and MGIT are summarized in **Table 4**.

**Table 4.** Diagnostic performance of urine mycobacterial LAM assay for tuberculous meningitis

| **Reference Standard** | **Sensitivity**<br>**n/N (%)** | **Specificity**<br>**n/N (%)** | **PPV**<br>**n/N (%)** | **NPV**<br>**n/N (%)** | **Positive**<br>**Likelihood**<br>**Ratio** | **Negative**<br>**Likelihood**<br>**Ratio** |
| --- | --- | --- | --- | --- | --- | --- |
| **Comprehensive Reference Standard** | 24/69<br>(34.8%) | 11/11<br>(100%) | 24/24<br>(100%) | 11/56<br>(19.6%) | - | 0.65 |
| **95% CI** | 23.7 to<br>47.2% | 71.5 to<br>100% | - | 17.1 to<br>22.5% | - | 0.6 to 0.7% |
| **Microbiological reference standard** | 14/23<br>(60.9%) | 47/57<br>(82.5%) | 14/24<br>(58.3%) | 47/56<br>(83.9%) | 3.47 | 0.47 |
| **95% CI** | 38.5 to<br>80.3% | 70.1 to<br>91.3% | 42.2 to<br>72.9% | 75.6 to<br>89.8% | 1.8 to 6.7% | 0.3 to 0.8% |
| **GX** | 12/19<br>(63.2%) | 49/61<br>(80.3%) | 12/24<br>(50%) | 49/56<br>(87.5%) | 3.2 | 0.5 |
| **95% CI** | 38.4 to<br>83.7% | 68.2 to<br>89.4% | 35.2 to<br>64.9% | 79.3 to<br>92.7% | 1.7 to 5.9% | 0.2 to 0.8% |
| **MGIT** | 8/12<br>(66.7%) | 52/68<br>(76.5%) | 8/24<br>(33.3%) | 52/56<br>(92.9%) | 2.83 | 0.44 |
| **95% CI** | 34.9 to<br>90.1% | 64.6 to<br>85.9% | 21.8 to<br>47.3% | 85.2 to<br>96.7% | 1.6 to 5.1% | 0.2 to 1.0% |

PPV-Positive predictive value; NPV-Negative predictive value; CI- Confidence Interval
GX-GeneXpert; MGIT- Mycobacterial Growth Indicator Tube

### Utility of Host Serum/ CSF Biomarkers in the Diagnosis of TBM

A comparison of six host biomarkers between TBM and non-TBM patients showed that serum IFN-γ and IL-1β levels were significantly different (*p* ≤ 0.05) between the two groups **(Table 5)**. ROC curve analysis **(Figure 2a-f)** demonstrated that serum IL-1β had an area under the ROC (AUC) of 0.943, with a sensitivity of 88.9%, and specificity of 90.9%. Serum IL-6 had an AUC of 0.442, with 28% sensitivity and 100% specificity, whereas serum IL-17A exhibited an AUC of 0.367, 20% sensitivity, and 100% specificity for TBM diagnosis.

**Figure 2.**
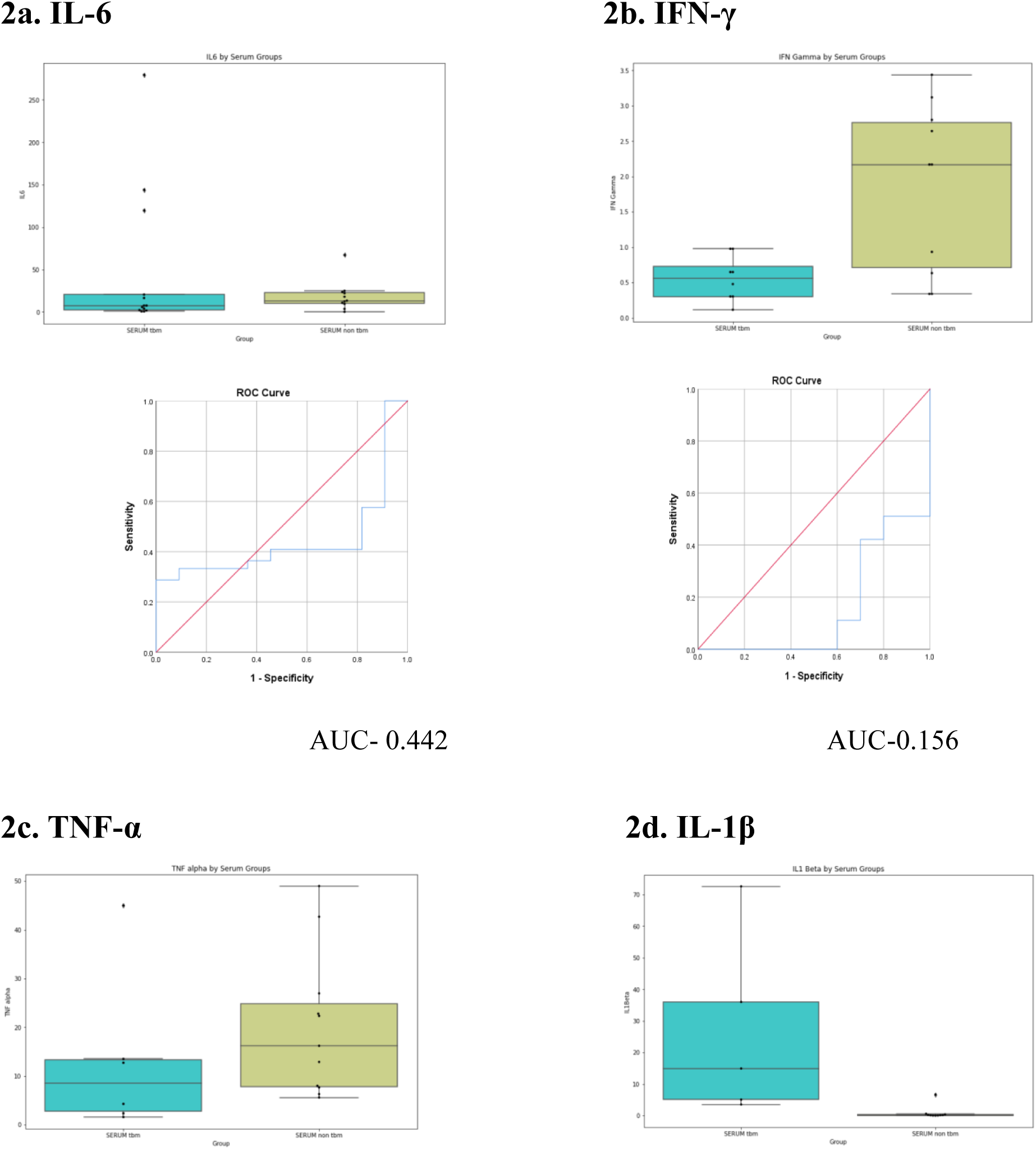
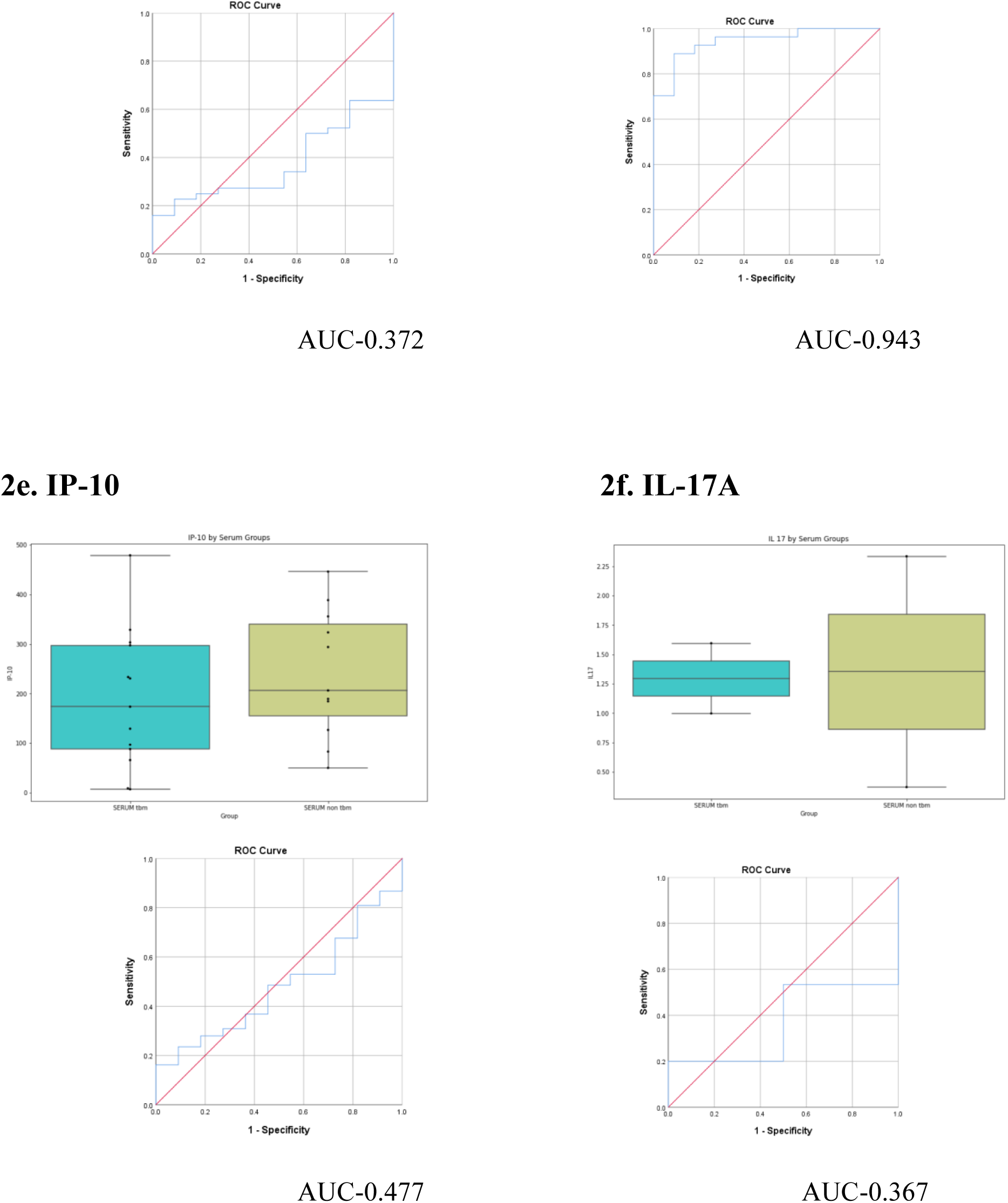
**(a-f). Boxplots showing levels of (a) IL-6, (b)IFN-γ, (c) TNF-α, (d) IL-1β, (e) IP-10, and (f) IL-17A in serum samples from TBM and non-TBM patients and ROC curves showing the accuracy of these biomarkers in the diagnosis of TBM.**

**Table 5.**
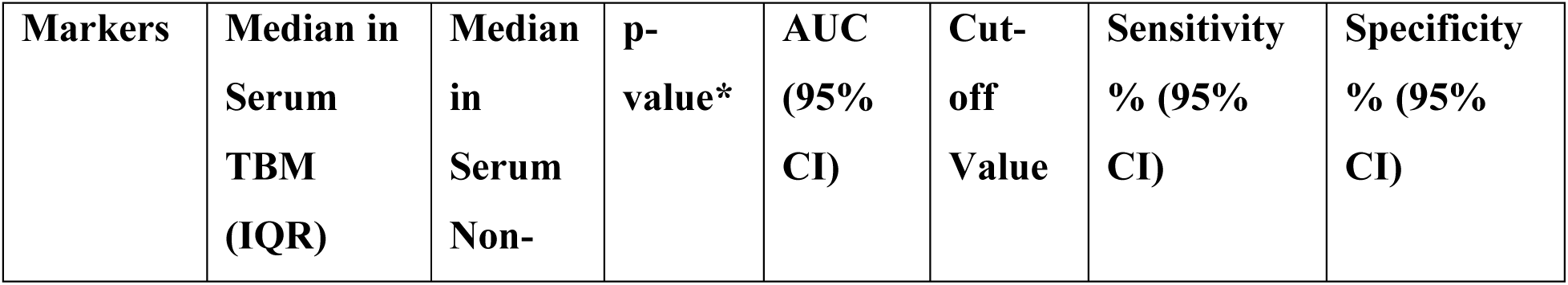
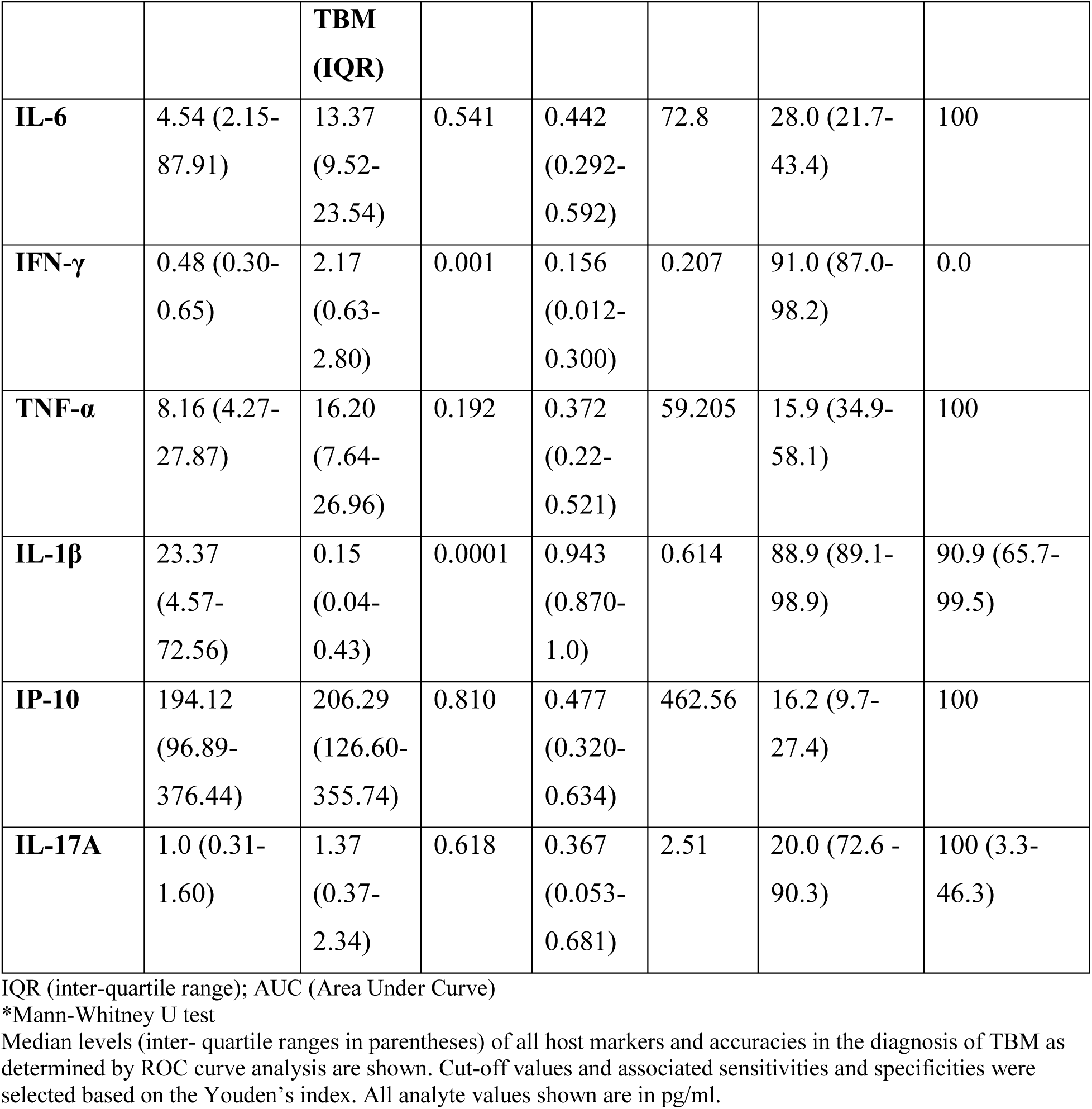
Utility of host biomarkers detectable in Serum samples from adults with suspected meningitis in the diagnosis of TB meningitis

| **Markers** | **Median in Serum TBM (IQR)** | **Median in Serum Non-** | **p-value\*** | **AUC (95% CI)** | **Cut-off Value** | **Sensitivity % (95% CI)** | **Specificity % (95% CI)** |
| --- | --- | --- | --- | --- | --- | --- | --- |

|  |  | **TBM**<br>**(IQR)** |  |  |  |  |  |
| --- | --- | --- | --- | --- | --- | --- | --- |
| **IL-6** | 4.54 (2.15-87.91) | 13.37<br>(9.52-23.54) | 0.541 | 0.442<br>(0.292-0.592) | 72.8 | 28.0 (21.7-43.4) | 100 |
| **IFN-γ** | 0.48 (0.30-0.65) | 2.17<br>(0.63-2.80) | 0.001 | 0.156<br>(0.012-0.300) | 0.207 | 91.0 (87.0-98.2) | 0.0 |
| **TNF-α** | 8.16 (4.27-27.87) | 16.20<br>(7.64-26.96) | 0.192 | 0.372<br>(0.22-0.521) | 59.205 | 15.9 (34.9-58.1) | 100 |
| **IL-1β** | 23.37<br>(4.57-72.56) | 0.15<br>(0.04-0.43) | 0.0001 | 0.943<br>(0.870-1.0) | 0.614 | 88.9 (89.1-98.9) | 90.9 (65.7-99.5) |
| **IP-10** | 194.12<br>(96.89-376.44) | 206.29<br>(126.60-355.74) | 0.810 | 0.477<br>(0.320-0.634) | 462.56 | 16.2 (9.7-27.4) | 100 |
| **IL-17A** | 1.0 (0.31-1.60) | 1.37<br>(0.37-2.34) | 0.618 | 0.367<br>(0.053-0.681) | 2.51 | 20.0 (72.6 -<br>90.3) | 100 (3.3-46.3) |

IQR (inter-quartile range); AUC (Area Under Curve)
\*Mann-Whitney U test
Median levels (inter- quartile ranges in parentheses) of all host markers and accuracies in the diagnosis of TBM as determined by ROC curve analysis are shown. Cut-off values and associated sensitivities and specificities were selected based on the Youden's index. All analyte values shown are in pg/ml.

ROC curve analysis of CSF biomarkers **(Figure 3a-f)** showed that IL-6, TNF-α, IL-1β, and IL-17 had AUC values exceeding 0.60. Among these, CSF IL-1β demonstrated the highest diagnostic accuracy, with an AUC of 0.750 (95% CI: 0.452-1.0), showing a sensitivity of 83.3% and a specificity of 71.4% **(Table 6).** CSF IL-6 and TNF-α exhibited similar sensitivities of 43.4% and 44.4%, and specificities of 81% and 100%, respectively.

**Figure 3.**
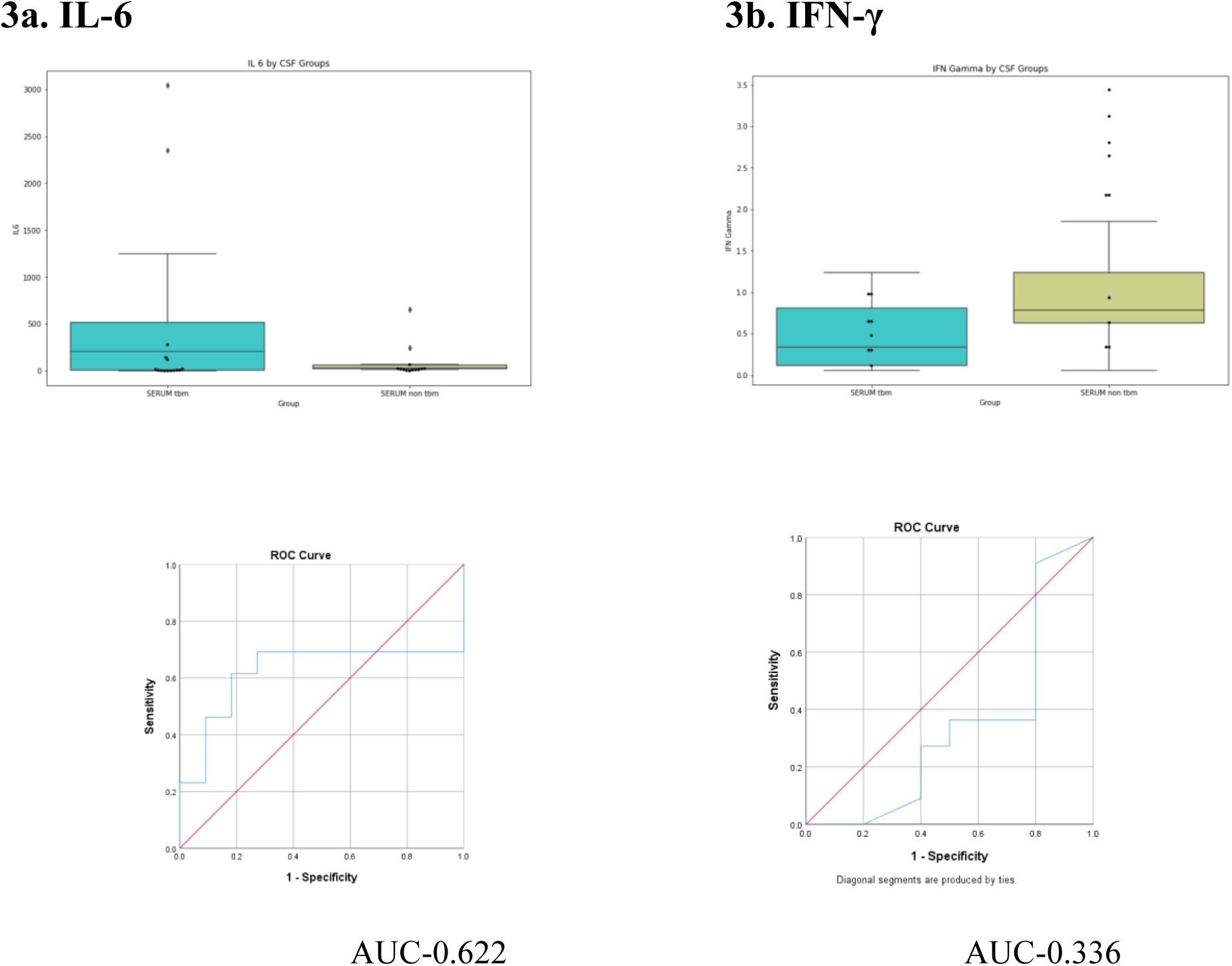
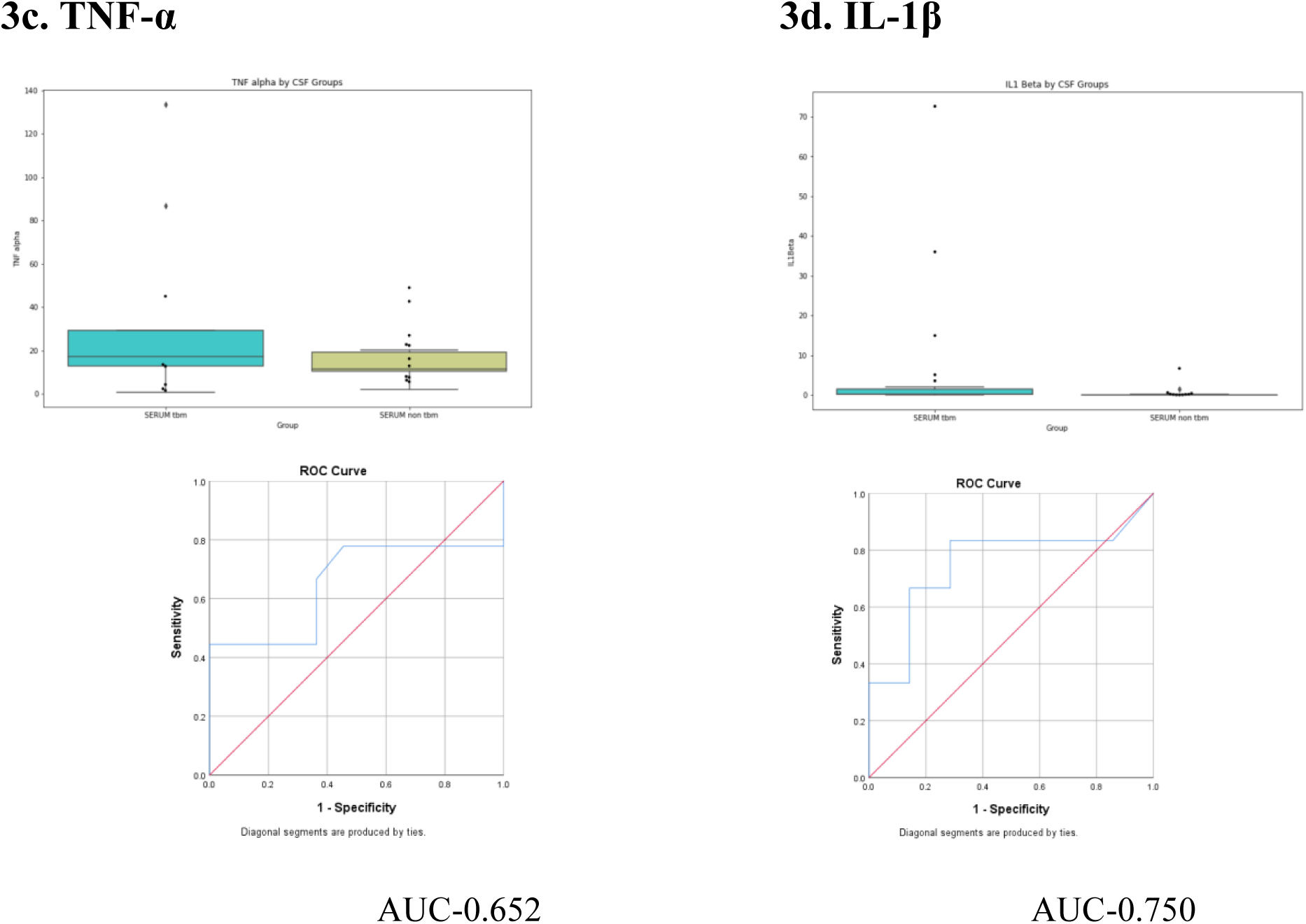
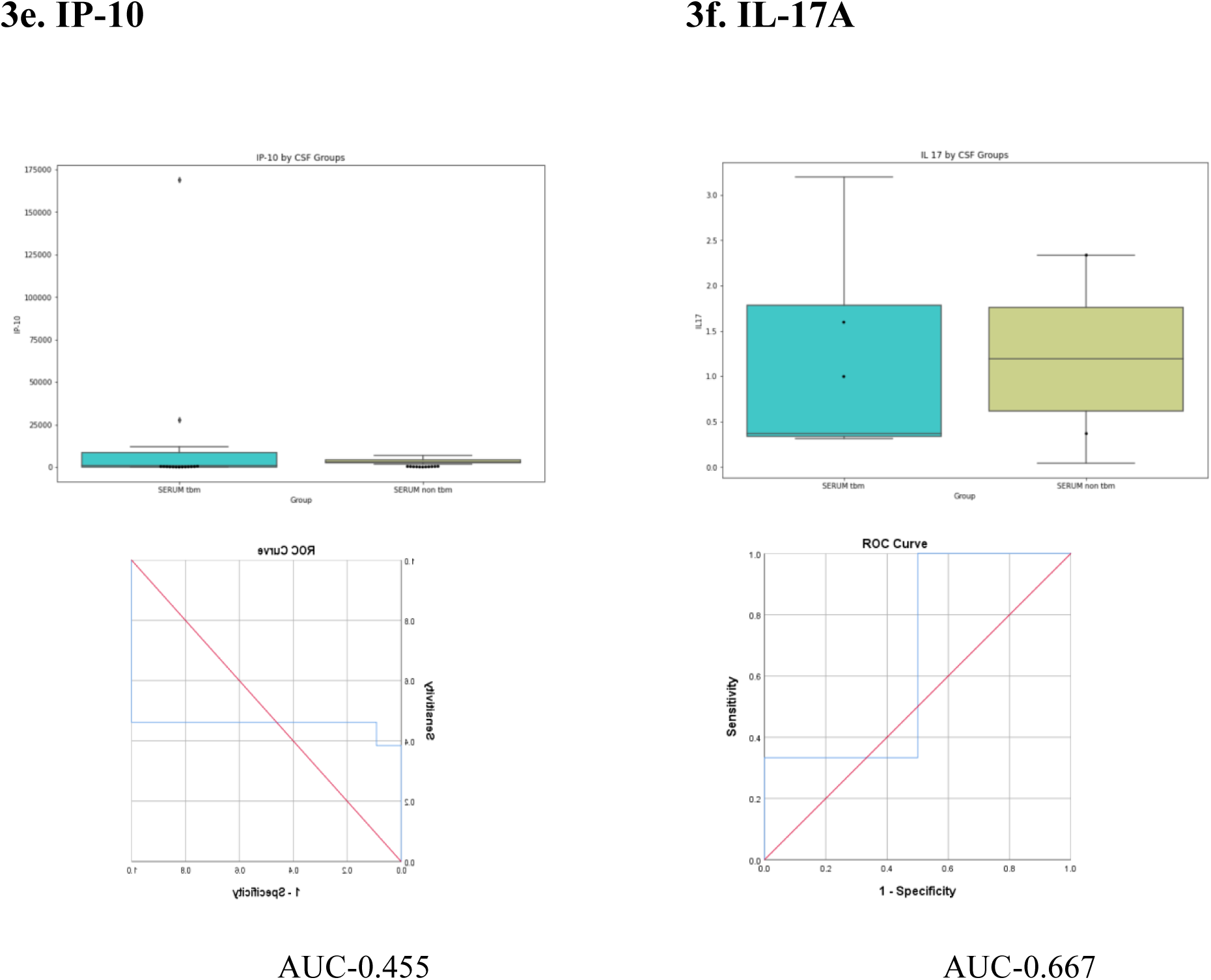
**(a-f). Boxplots showing levels of (a) IL-6, (b)IFN-γ, (c) TNF-α, (d) IL-1β, (e) IP-10, and (f) IL-17A in CSF samples from TBM and non-TBM patients and ROC curves showing the accuracy of these biomarkers in the diagnosis of TBM.**

**Table 6.**
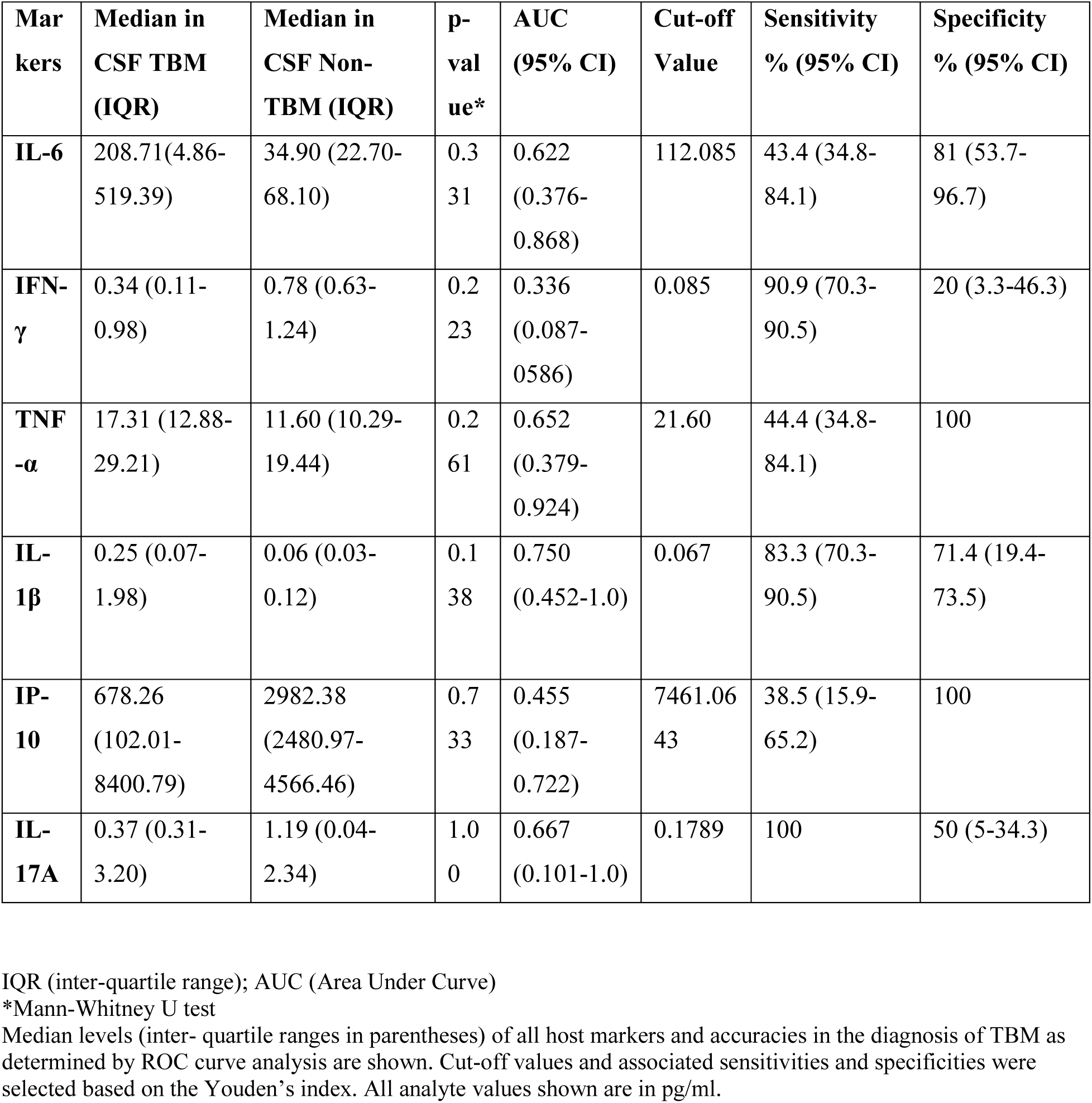
Utility of host biomarkers detectable in CSF samples from adults with suspected meningitis in the diagnosis of TB meningitis

| **Mar**<br>**kers** | **Median in**<br>**CSF TBM**<br>**(IQR)** | **Median in**<br>**CSF Non-**<br>**TBM (IQR)** | **p-**<br>**val**<br>**ue\*** | **AUC**<br>**(95% CI)** | **Cut-off**<br>**Value** | **Sensitivity**<br>**% (95% CI)** | **Specificity**<br>**% (95% CI)** |
| --- | --- | --- | --- | --- | --- | --- | --- |
| **IL-6** | 208.71(4.86-519.39) | 34.90 (22.70-68.10) | 0.3<br>31 | 0.622<br>(0.376-0.868) | 112.085 | 43.4 (34.8-84.1) | 81 (53.7-96.7) |
| **IFN-**<br>**γ** | 0.34 (0.11-0.98) | 0.78 (0.63-1.24) | 0.2<br>23 | 0.336<br>(0.087-0.586) | 0.085 | 90.9 (70.3-90.5) | 20 (3.3-46.3) |
| **TNF-**<br>**α** | 17.31 (12.88-29.21) | 11.60 (10.29-19.44) | 0.2<br>61 | 0.652<br>(0.379-0.924) | 21.60 | 44.4 (34.8-84.1) | 100 |
| **IL-**<br>**1β** | 0.25 (0.07-1.98) | 0.06 (0.03-0.12) | 0.1<br>38 | 0.750<br>(0.452-1.0) | 0.067 | 83.3 (70.3-90.5) | 71.4 (19.4-73.5) |
| **IP-**<br>**10** | 678.26<br>(102.01-8400.79) | 2982.38<br>(2480.97-4566.46) | 0.7<br>33 | 0.455<br>(0.187-0.722) | 7461.06<br>43 | 38.5 (15.9-65.2) | 100 |
| **IL-**<br>**17A** | 0.37 (0.31-3.20) | 1.19 (0.04-2.34) | 1.0<br>0 | 0.667<br>(0.101-1.0) | 0.1789 | 100 | 50 (5-34.3) |

IQR (inter-quartile range); AUC (Area Under Curve)
\*Mann-Whitney U test
Median levels (inter- quartile ranges in parentheses) of all host markers and accuracies in the diagnosis of TBM as determined by ROC curve analysis are shown. Cut-off values and associated sensitivities and specificities were selected based on the Youden's index. All analyte values shown are in pg/ml.

### Utility of Host Serum/CSF Biomarkers in the Prognosis of TBM

Serum biomarker levels at baseline and FU-1 (one month after starting treatment) were measured in all TBM patients, excluding four who died before follow-up. Biomarker levels were analyzed in relation to treatment outcomes: early responders (n=2), late responders (n=9), and on-treatment mortality (n=13). The trend in serum biomarkers was analyzed in patients (n=9) with high baseline serum IP-10 levels (>400 pg/ml). Additionally, serum and CSF biomarker concentrations were compared between TBM (serum: n=69; CSF: n=13) and non-TBM patients (serum and CSF: n=11). CSF biomarker levels were also evaluated in relation to mortality (n=13).

Among the TBM patients who died, serum and CSF IP-10 levels were analyzed at baseline (n=13) and at FU-1 (n=9). Three of the four patients who died before FU-1 had elevated baseline serum ( > 200 pg/ml) and CSF IP-10 levels (> 8000 pg/ml). Baseline CSF IP-10 concentrations were markedly higher than serum levels (median=6447.79 vs 164.59 pg/ml). Rising serum and CSF IP-10 levels at FU-1 were observed, with elevated serum levels in 8 patients and increased CSF levels in 7 patients **(Figure 4a–i)**. Two patients with initially very high CSF IP-10 levels **(Figure 5a and 5c)** later died (at third and fifth months of treatment) despite slight declines at one month.

**Figure 4.**
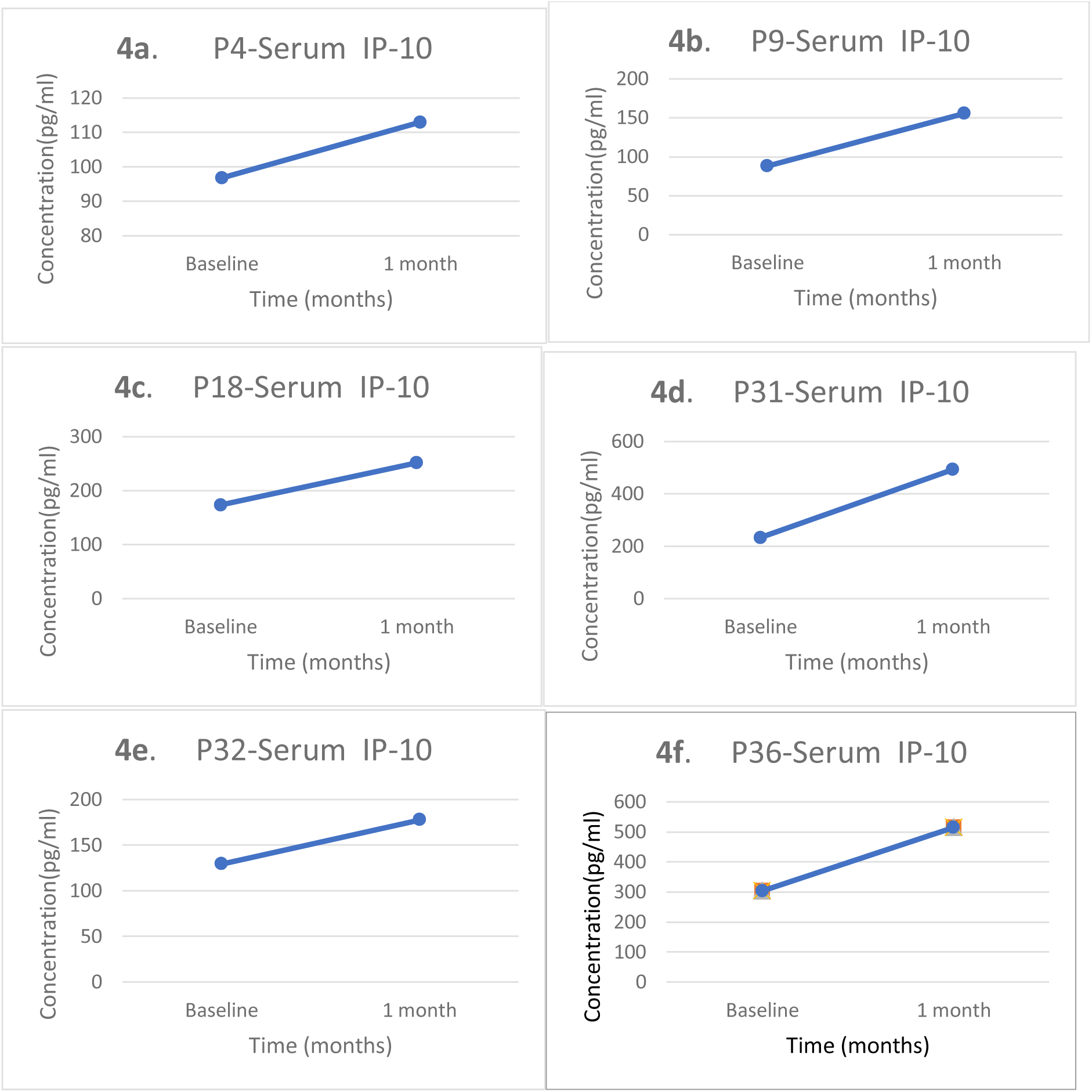
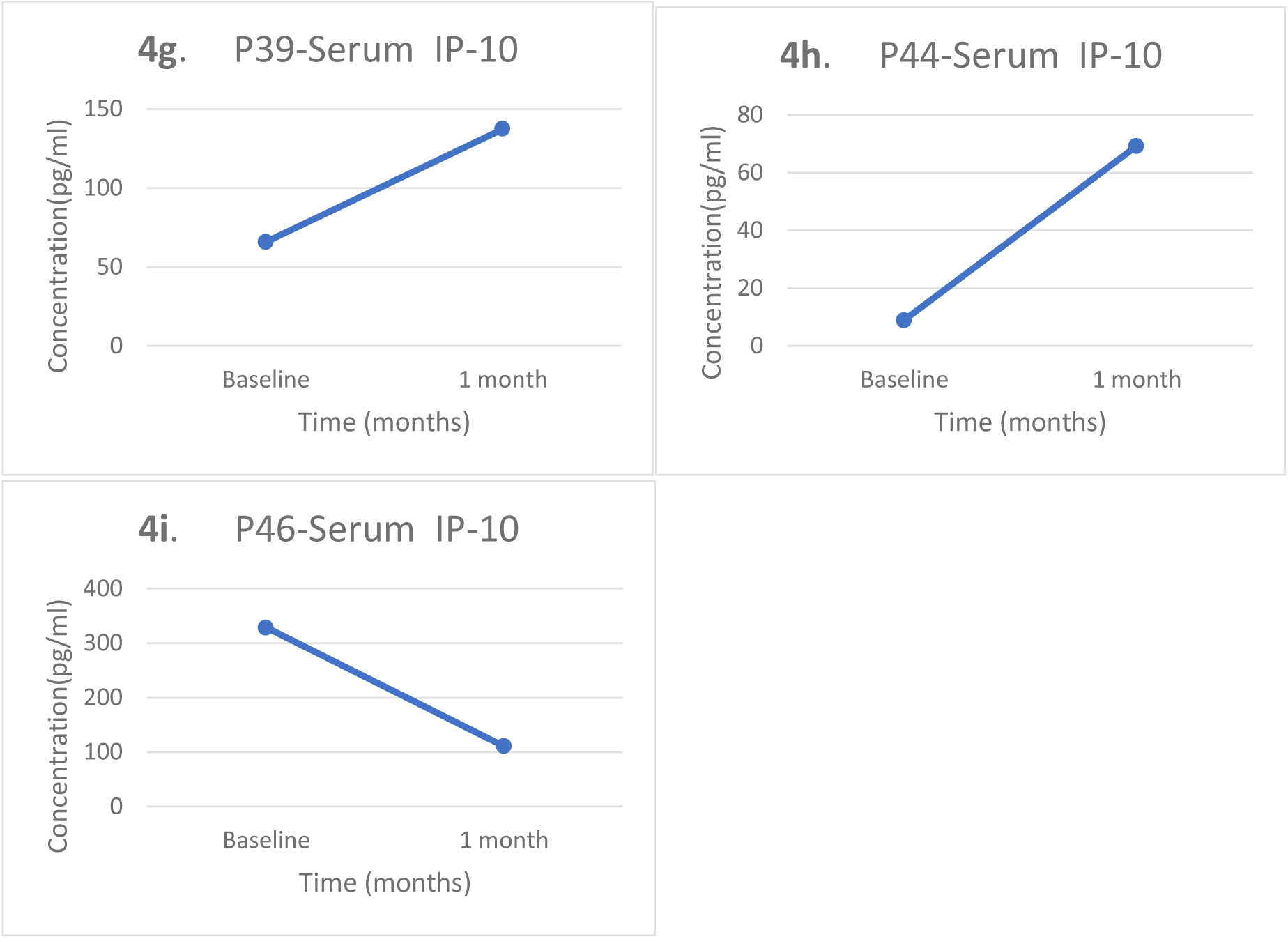
(a–i) shows the trend in serum IP-10 at baseline and 1 monthly follow-up among the TBM patients. Concentrations of all analytes shown are in pg/ml.

**Figure 5.**
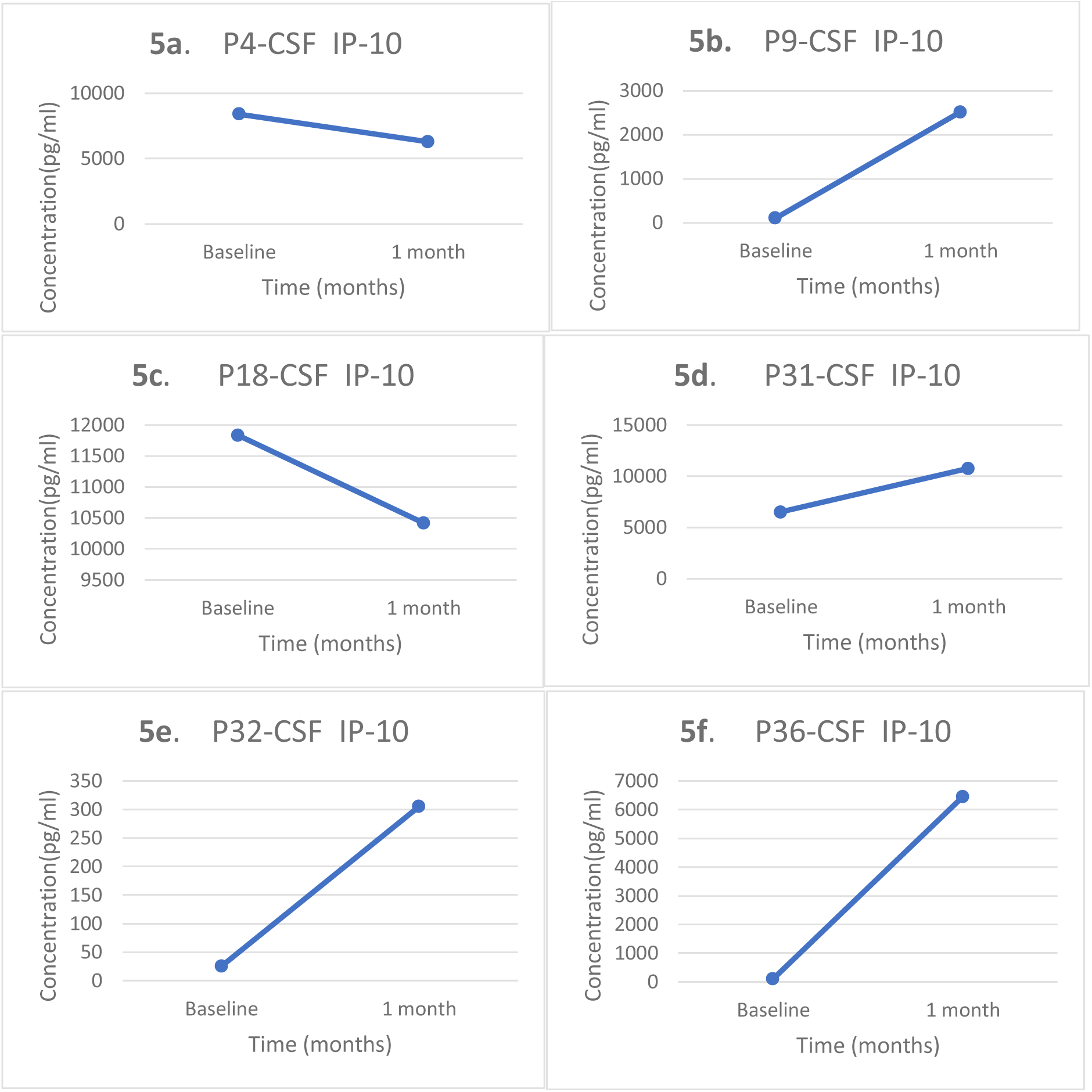
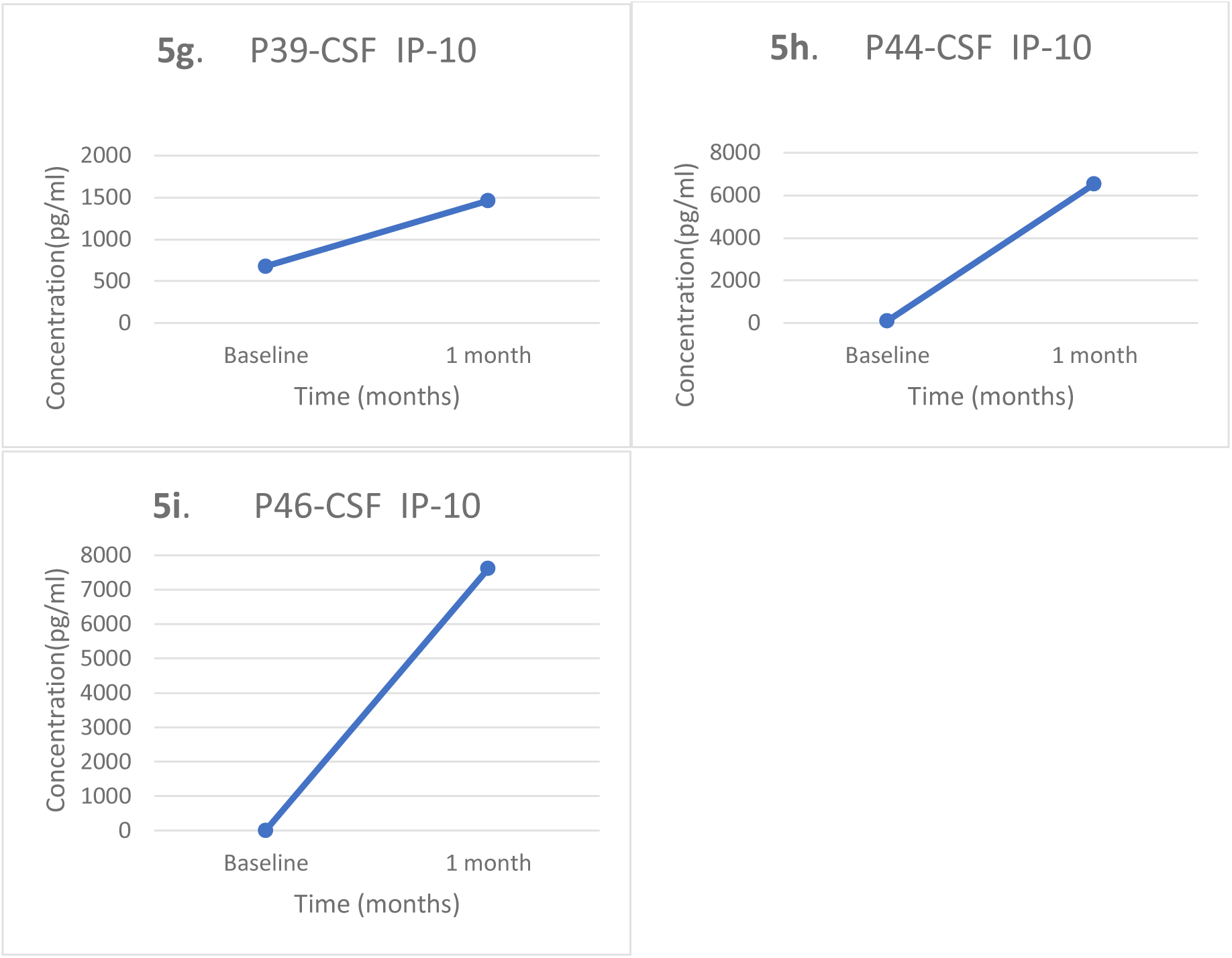
(a–i) shows the trend in serum IP-10 at baseline and 1 monthly follow-up among the TBM patients. Concentrations of all analytes shown are in pg/ml.

Serum TNF-α and IL-6 levels decreased within one month in the early responder group (n=2) **(Figures 6a and 6b).** Conversely, serum IP-10 levels increased slightly in the early responder group (n=2) **(Figure 6c)**. Patients with a high baseline serum IP-10 level exceeding 400 pg/ml (n=13) were also monitored for changes in biomarker levels after one month **(Figure 7).** At the follow-up, eight patients showed a decreasing trend in serum IP-10 levels, consistent with their clinical improvement. Conversely, five individuals with delayed clinical improvement exhibited higher IP-10 levels at follow-up, suggesting that serum IP-10 may decrease more gradually than other inflammatory markers.

**Figure 6.**
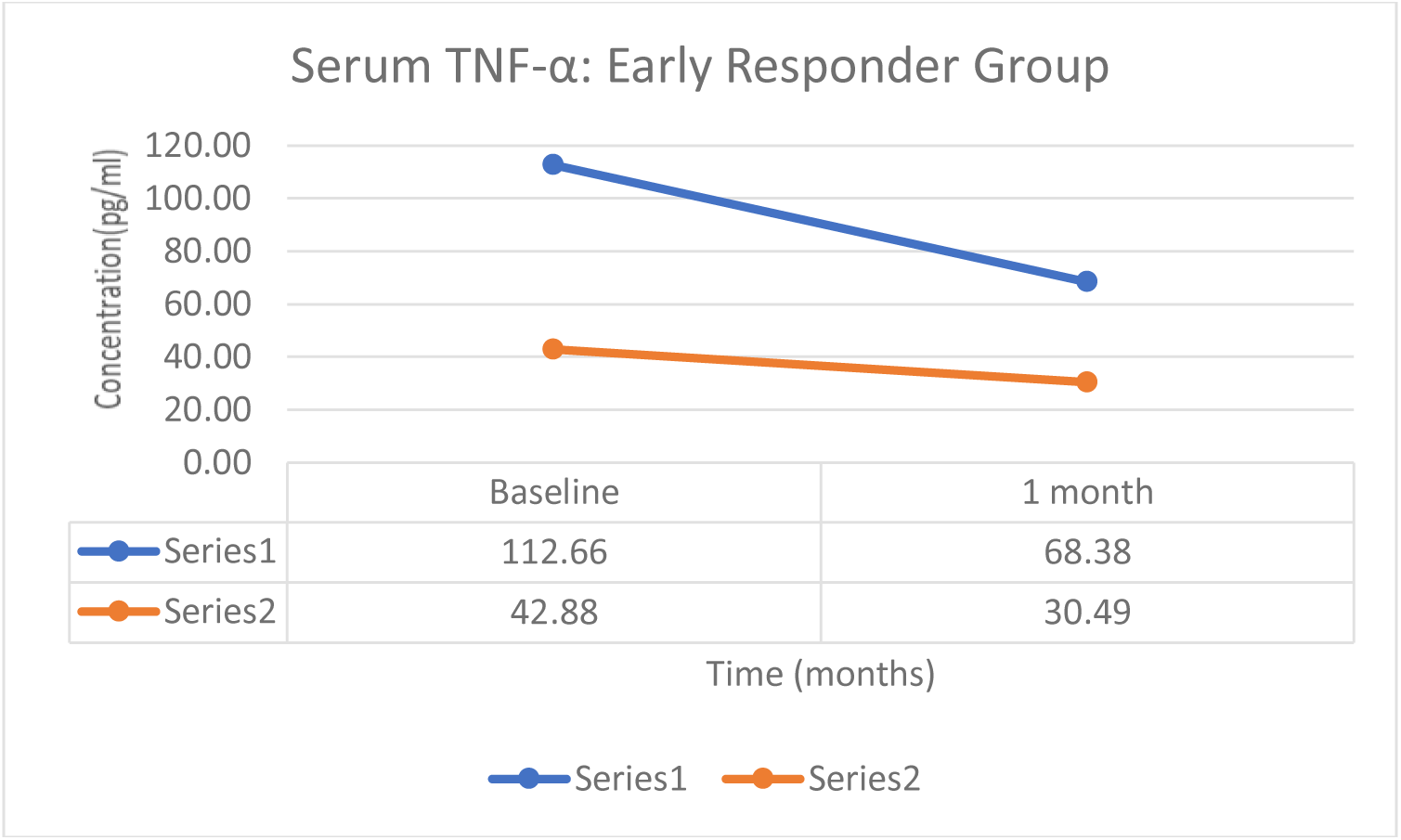
**shows the trend in Serum TNF-α at baseline and at 1-month follow-up among TBM patients (early responder group).** Concentrations of all analytes shown are in pg/ml.

**Figure 6b.**
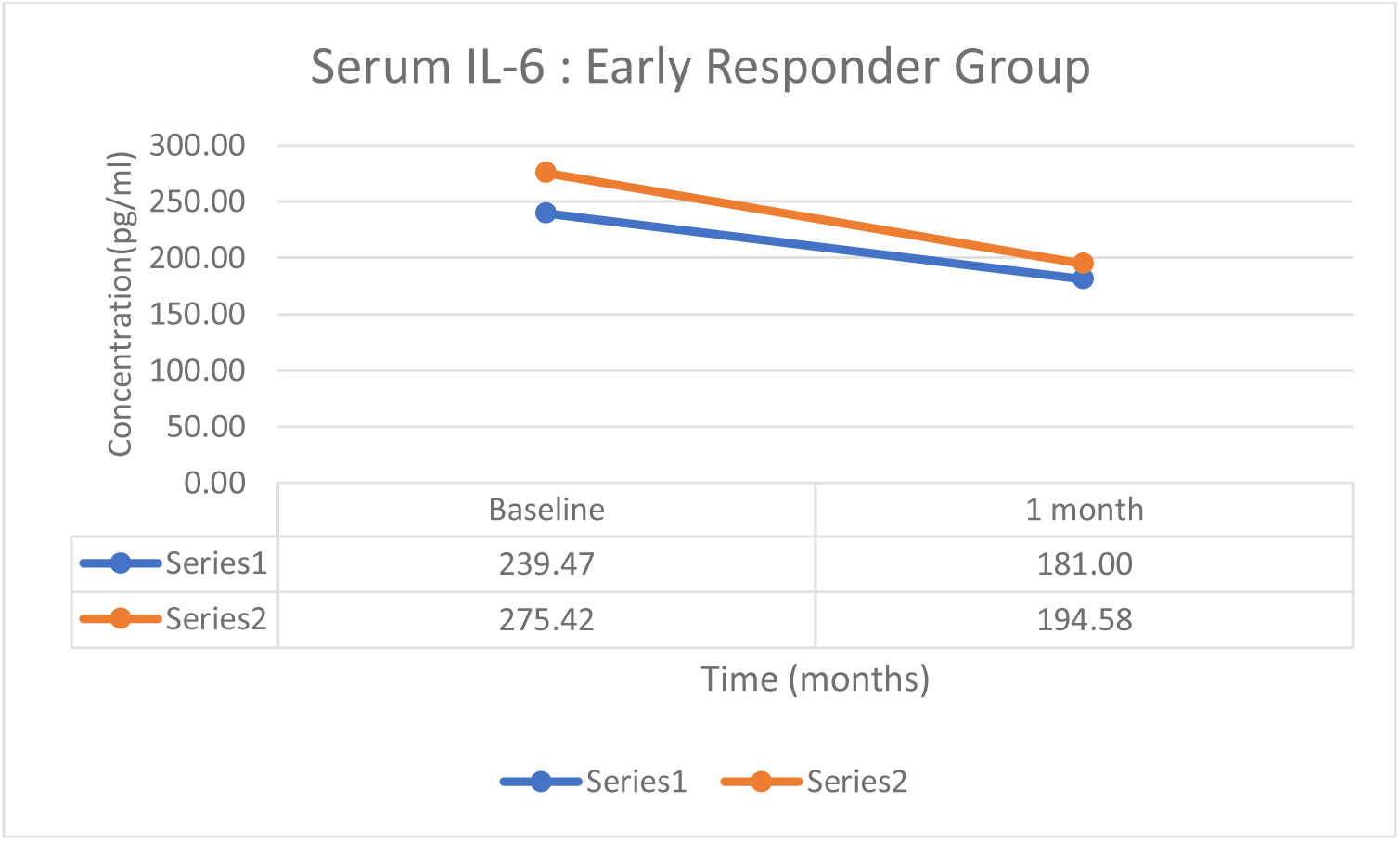
**shows the trend in Serum IL-6 at baseline and at 1-month follow-up among TBM patients (early responder group).** Concentrations of all analytes shown are in pg/ml.

**Figure 6c.**
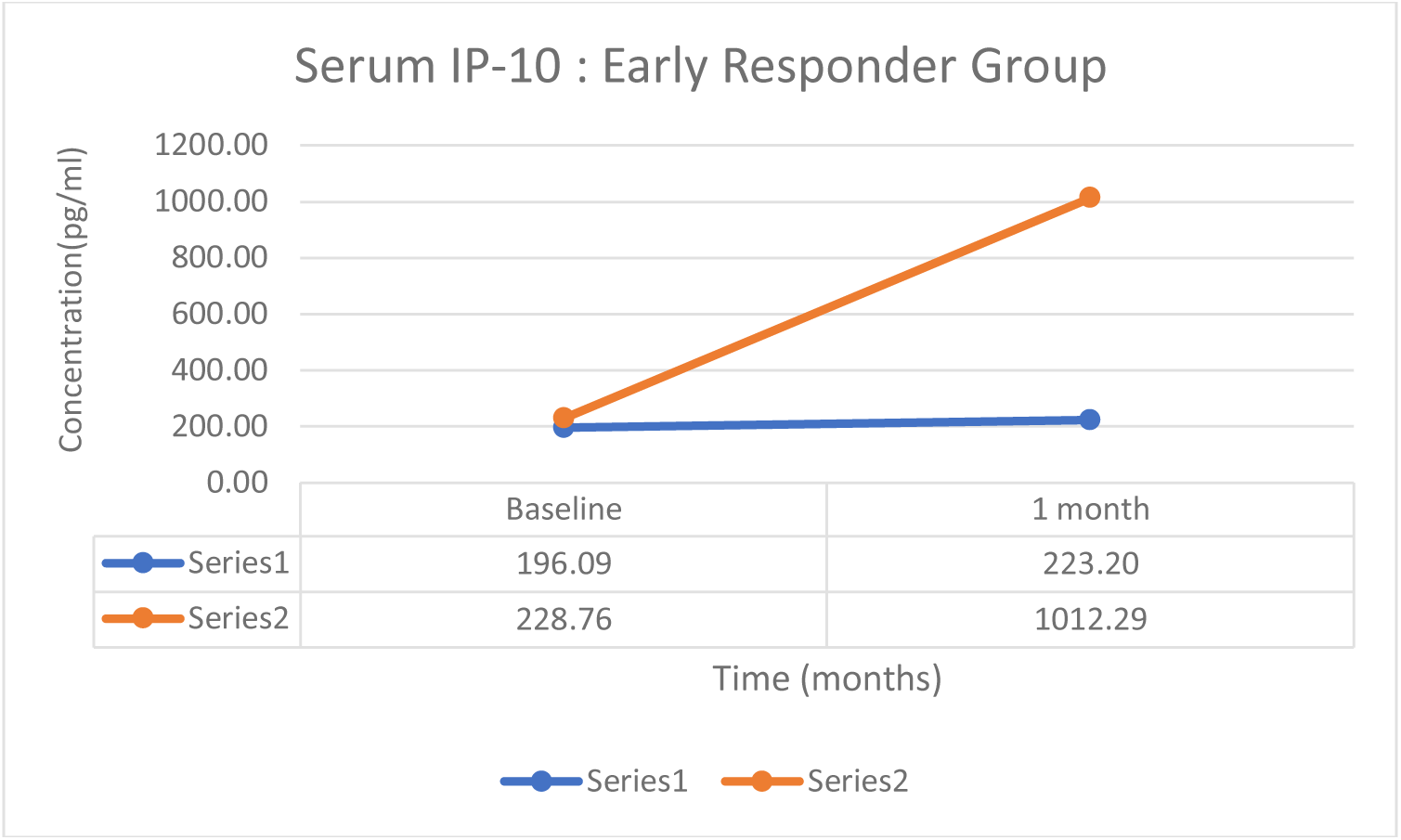
**shows the trend in Serum IP-10 at baseline and at 1-month follow-up among TBM patients (early responder group).** Concentrations of all analytes shown are in pg/ml.

**Figure 7.**
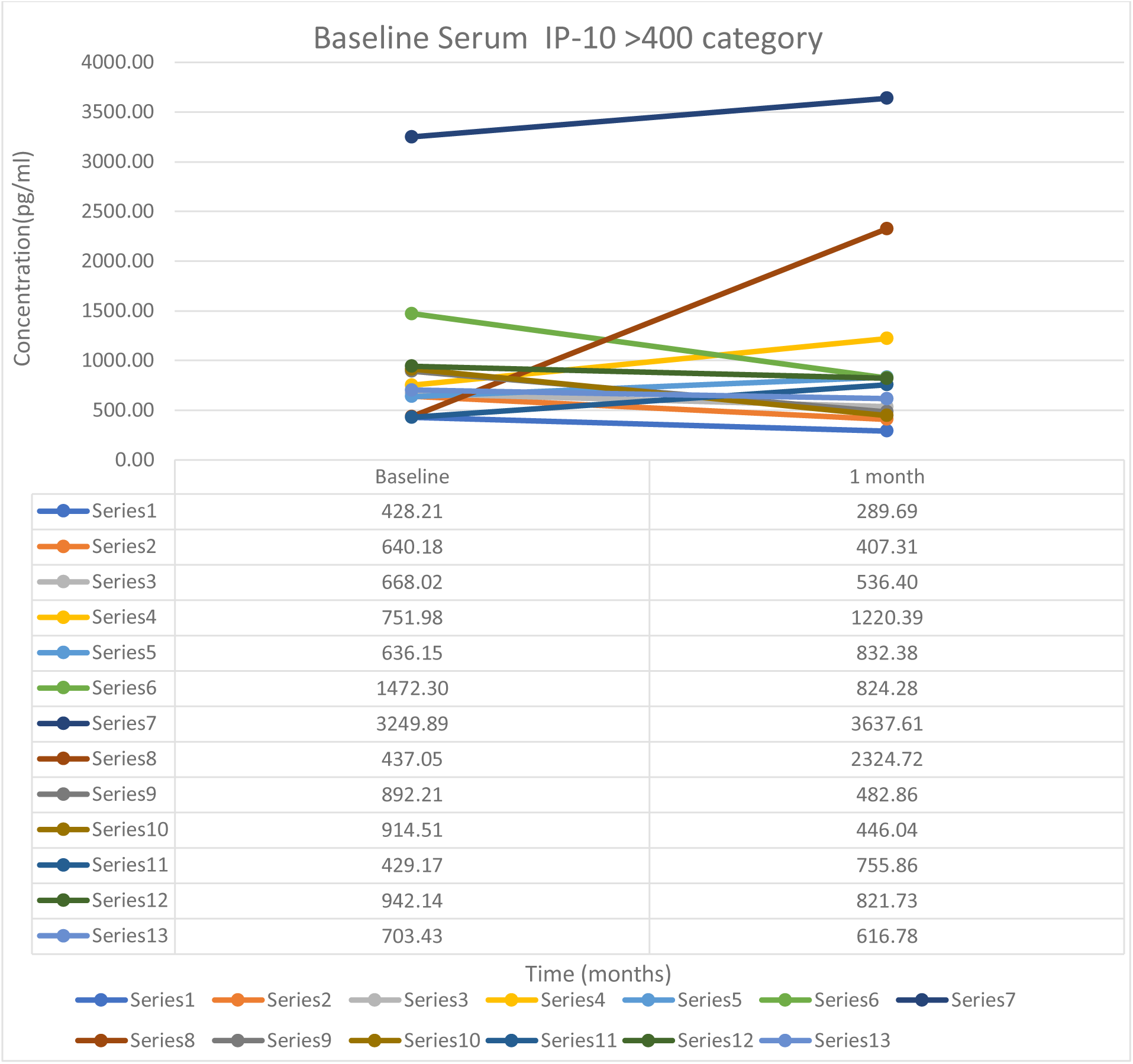
shows the trend in Serum IP-10 at baseline and at 1-month follow-up among TBM patients (Baseline serum IP-10> 400 pg/ml; n=13). Concentrations of all analytes shown are in pg/ml.

Among late responders (n=9), six patients had rising serum IP-10 at FU-1, while three showed a slight decline **(Figure 8; series 5, 6, 9).** Although one of the three patients had begun clinically responding to treatment after one month, symptoms persisted for a longer period of time, necessitating an extension of treatment for 15 months.

**Figure 8.**
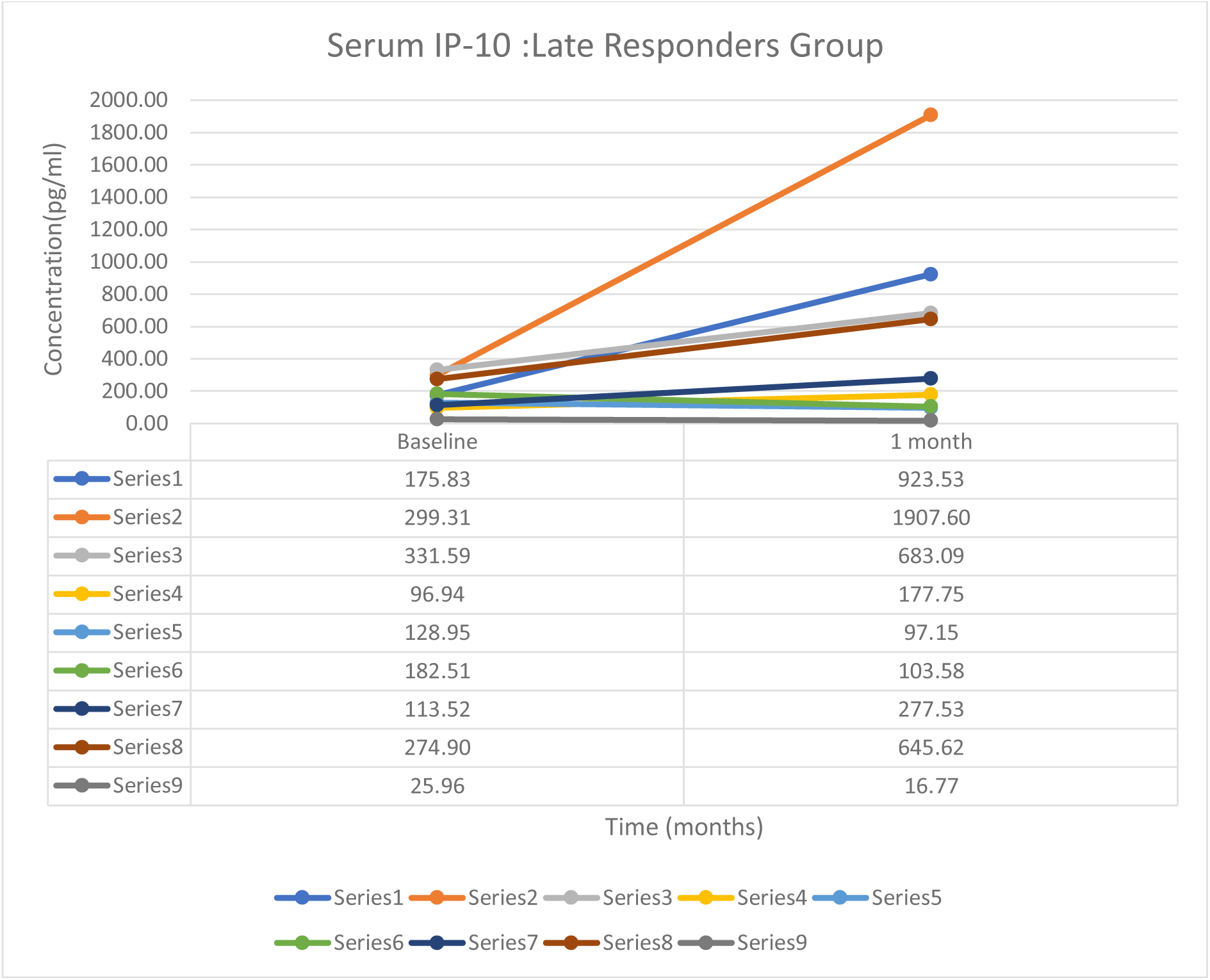
shows the trend in Serum IP-10 at baseline and at 1-month follow-up among TBM patients (late responder group). Concentrations of all analytes shown are in pg/ml.

## Discussion

Current estimates keep the number of undiagnosed cases at 2.9 million of the yearly 10 million new tuberculosis cases. (13) TBM diagnosis is difficult due to the low sensitivity of microscopy, prolonged culture turnaround time, and limited sensitivity and high cost of molecular assays such as GX. Consequently, TBM is often diagnosed using a combination of clinical, laboratory, and radiological findings. (14)

Most previous research has focused on detecting LAM antigen in urine in HIV-infected patients and children with TBM (23,24). Our earlier research showed that urinary LAM antigen testing demonstrated high sensitivity and specificity in diagnosing paediatric tuberculosis, leading to a 38.5% increase in case detection for intrathoracic TB and a 41.6% increase for lymph node TB (17).The current study aimed to evaluate the effectiveness of urine and CSF LAM antigen testing for diagnosing TB meningitis in immunocompetent adults.

CSF LAM LFA showed moderate sensitivity (43.5%) and specificity (80.7%) compared with MRS. When compared with CRS, sensitivity decreased to 30.4%, while specificity increased to 100%. These findings are similar to those of *Kwizera et al.* (18), who reported 33% sensitivity and 96% specificity for definite TBM, which fell to 24% and 95% for definite/probable TBM (18).

In our study, CSF smear microscopy failed to detect any TBM cases, whereas CSF LAM identified 10 of 23 definite TBM cases, demonstrating a superior diagnostic yield over conventional smear methods. The low MGIT culture yield may be attributable to most patients already receiving treatment prior to CSF sampling. False-negative CSF LAM test results in definite TBM may be related to the paucibacillary nature of TBM, low CSF LAM concentrations, and sequestration of LAM within immune complexes. (19)

We identified 11 patients who met the criteria for probable TBM (GX/MGIT-negative) and had positive CSF LAM LFA results. This suggests that LAM and PCR-based assays may detect different patient subsets. One explanation is that LAM detects cell-wall antigens in CSF even when intact bacilli or nucleic acids are insufficient for molecular or culture detection (20). The association between CSF LAM positivity, TBM classification as ‘probable’, and high mortality supports the conclusion that these results are likely true positives. (20)

Urine LAM LFA demonstrated a higher sensitivity of 60.9% against MRS compared to CSF LAM at 43.5%. Against CRS, the test showed low sensitivity (30.4%) but perfect specificity (100%). These results align with previous meta-analyses, which reported pooled sensitivities of 31–42% and specificities of 91–97% (21–23). Overall, urine LAM exhibited greater sensitivity than CSF LAM while maintaining similar specificity. Higher sensitivity in urine can be ascribed to renal filtration and urinary excretion of circulating LAM antigen (24).

High specificity and PPV support the use of LAM as a rule-in test within diagnostic algorithms. Reduced specificity reported in some studies may result from imperfect reference standards, cross-reactivity with urinary pathogens, or non-tuberculous mycobacteria. (25). Overall, these findings suggest that the TB-LAM LFA can be used as an initial, routine point-of-care rule-in test for TBM, particularly in high TB-burden settings.

### Utility of Host Serum/ CSF Biomarkers in the Diagnosis and Prognosis of TBM

Among the CSF biomarkers evaluated, IL-1β demonstrated the highest diagnostic accuracy, with a sensitivity of 83.3% and specificity of 71.4% for diagnosing TBM. *Koeken et al.* reported that CSF IL-1β levels were 44-fold higher in TBM patients than in controls (26). Significant differences (p ≤ 0.05) in serum IL-1β levels between TBM patients and controls suggest a distinct inflammatory cytokine profile that may help differentiate TBM from other CNS infections, such as cryptococcal meningitis.

CSF IL-6, TNF-α, and IP-10 demonstrated moderate sensitivity with high specificity, whereas CSF IL-17A exhibited 100% sensitivity but modest specificity (50%). *Amene Saghazadeh et al.* similarly reported that CSF levels of IFN-γ, IL-1β, TNF-α, IL-6, and IL-17 were significantly higher in TBM than in cryptococcal, viral, or aseptic meningitis, emphasizing their role in CNS inflammation and immune activation in TBM. (27).

Among serum biomarkers, IL-1β showed the highest diagnostic accuracy, with 88.9% sensitivity and 90.9% specificity, while serum IL-6 and IL-17A showed moderate sensitivity and excellent specificity. This supports the potential utility of combined serum biomarkers as non-invasive screening tools for TBM (28). Previous research by *Alhan et al*. also found notably higher serum and CSF IL-6 levels in TBM patients compared to controls, confirming IL-6 as a valuable biomarker for early diagnosis. (29).

Furthermore, serum and CSF biomarkers showed important prognostic value in TBM. Patients who died within one month had markedly elevated baseline IP-10 levels in serum and CSF, while the remaining fatal cases showed a rising trend. These observations suggest that elevated IP-10 levels are associated with poor outcomes, and patients should be closely monitored. Similarly, findings from *Simona et al*. demonstrated that consistently high IP-10 levels were associated with poor treatment response (30).

Serum IL-6 and TNF-α levels decreased in patients who responded rapidly to treatment, suggesting their potential as markers of early therapeutic response. *Misra et al.* observed that TBM patients exhibited high CSF IL-6 levels, which declined after antitubercular therapy, indicating its role as a marker of inflammation and treatment effectiveness. (31).

Overall, IL-1β demonstrated high sensitivity and specificity for TBM diagnosis in both CSF and serum, while elevated IP-10 levels were associated with an adverse prognosis. Research by *Sinha et al*. has also linked elevated CSF and serum TNF-α and IL-1β levels to severe neuroinflammation, advanced TBM, tuberculoma formation, and poor clinical outcomes (32).

Collectively, these findings indicate that using a combined cytokine panel incorporating IL-1β, IL-6, TNF-α, IL-17A, and IP-10 could enhance diagnostic accuracy and therapeutic monitoring.

The present study has multiple strengths. Its prospective design minimized selection bias and enabled standardized clinical and laboratory assessment. Use of consensus TBM case definitions ensured uniform patient classification and improved the reliability of diagnostic accuracy estimates. Patients were systematically evaluated, enabling development of a CRS for evaluating the performance of urine and CSF LAM. Furthermore, LAM detection was assessed in both CSF and urine samples from the same cohort, permitting direct comparison of diagnostic utility across specimen types and reduced intra-study variability.

## Conclusion

In conclusion, LAM antigen is a highly specific pathogen biomarker for diagnosing TBM. LAM showed high specificity with moderate sensitivity in both urine and CSF, supporting its use as a valuable rule-in test for TBM. Its superior performance in urine highlights the utility of a non-invasive bedside diagnostic sample.

Among host biomarkers, IL-1β showed the highest diagnostic performance in both serum and CSF, supporting its role as a promising marker of CNS inflammation and disease activity in TBM. Elevated levels of IFN-γ, IL-6, TNF-α, and IL-17A further indicate the immune activation associated with TBM and can help differentiate it from other CNS infections.

Several host biomarkers also demonstrated prognostic utility. Consistently elevated IP-10 levels correlated with worse clinical outcomes and higher mortality, indicating its use as a marker for disease severity. Similarly, declining serum IL-6 and TNF-α levels after therapy were useful markers for monitoring treatment response.

Overall, these findings indicate that a combined approach incorporating pathogen-specific biomarkers, such as LAM antigen, and host inflammatory cytokines, including IL-1β, IL-6, TNF-α, IL-17A, IFN-γ, and IP-10, could enhance early diagnosis, prognosis, and treatment monitoring of TBM.

## Supplementary Datasets

**Supplementary 1.** Study design and diagnostic algorithm for TB-LAM detection in CSF and urine

**Supplementary 2.** CT findings in study groups

**Supplementary 3a.**CSF parameters among study groups

**Supplementary 3b.** Other CSF parameters among the study groups

**Supplementary 4.** BMRC staging of study groups

**Supplementary 5.** Clinical outcome of study participants

## Author’s contributions

UBS conceptualized the study and planned the work. UBS and AKP performed the literature search and drafted the manuscript. UBS, AKP, and AAK contributed to diagnostics and formal analysis. UK, and KS contributed to laboratory management. UBS and AKP contributed to study design, clinical discussion, and data analysis. AAK, NW, and AKS contributed to clinical data collection and patient care.

## Availability of data and materials

The datasets generated during and/or analyzed during the current study are available from the corresponding author on reasonable request.

## Ethical approval

Ethical approval was obtained from the Institutional Ethics Committee, AIIMS, New Delhi (Letter No. IECPG-59/27.01.2021).

## Patient Consent Statement

The study was conducted in accordance with the ethical principles of the Declaration of Helsinki of the World Medical Association. Participants or their legal guardians provided written informed consent prior to enrolment. All data were anonymized ahead of analysis to ensure confidentiality.

## Consent to publish

Patients or their parents/guardians signed informed consent to the publication of their data.

## Acknowledgments

The work reported was the analysis of diagnostic tests, carried out to support patient care.

